# Donor strand complementation, isopeptide bonds and glycosylation reinforce highly resilient archaeal thread filaments

**DOI:** 10.1101/2022.04.26.489512

**Authors:** Matthew C. Gaines, Michail N. Isupov, Shamphavi Sivabalasarma, Risat Ul Haque, Mathew McLaren, Patrick Tripp, Alexander Neuhaus, Vicki Gold, Sonja-Verena Albers, Bertram Daum

**Affiliations:** Living Systems Institute, University of Exeter, Stocker Road, EX4 4QD, Exeter, United Kingdom; Department of Biosciences, College of Life and Environmental Sciences, Stocker Road, EX4 4QD, Exeter, United Kingdom; Henry Wellcome Building for Biocatalysis, Biosciences, College of Life and Environmental Sciences, University of Exeter, EX4 4QD, Exeter, United Kingdom; Institute of Biology II, Molecular Biology of Archaea, University of Freiburg, Schänzlestraße 1, 79104 Freiburg, Germany; Spemann Graduate School of Biology and Medicine, University of Freiburg, Freiburg, Germany; Signalling Research Centres BIOSS and CIBBS, Faculty of Biology, University of Freiburg, Freiburg, Germany

## Abstract

Pili are ubiquitous filamentous surface extensions that play crucial roles for bacterial and archaeal cellular processes such as adhesion, biofilm formation, motility, cell-cell communication, DNA uptake and horizontal gene transfer to name a few. Here we report on the discovery and structure of the archaeal thread – a remarkably stable archaeal pilus that belongs to a so-far largely unknown class of protein filaments. We find that the filament is highly glycosylated and interconnected via donor strand complementation, as well as isopeptide bonds, reminiscent of bacterial type I pili. Despite striking structural similarity with bacterial type-1 pili, archaeal threads appear to have evolved independently and are likely assembled by a markedly distinct mechanism.

## Introduction

Prokaryotes assemble various cells surface structures with a plethora of functions. Besides the flagellum that is crucial for bacterial motility, bacteria also assemble numerous types of pili ranging from type I pili (T1P) to type V pili (T5P), where each is filament type fulfils a variety of specific tasks. Regardless, all these types of pili have in common that membrane spanning multiprotein complexes aid their assembly and anchor the filaments in the cell envelope ^1, 2^. Like bacteria, most archaea possess numerous cell surface filaments ^3–5^. The most extensively investigated archaeal surface filaments belong to the type IV filament superfamily (T4FF). Co-option and adaptation of homologous genes in the T4FF resulted in a variety of Type-IV pili (T4P) - like filaments with functions ranging from adhesion and motility to DNA exchange ^4, 6^.

The crenarchaeon *S. acidocaldarius* serves as an intriguing model organism to study cell surface structures, as it thrives at 75 °C and pH 3 and hence surface structures are adapted to these extreme environmental conditions^7^. One of the best studied surface structures in *S. acidocaldarius* is the archaellum, which is a gyrating T4FF that enables cells to swim through liquid media and is found in many motile archaea^6, 8, 9^. The filament, which is formed by signal peptide (SPIII)-cleaved and N-terminally processed pre-archaellins, has a helical symmetry and is highly glycosylated ^10–13^. Four structures of archaella were solved by CryoEM to date^12, 14–16^. All these structures revealed that the archaellins are tadpole-shaped monomers with a hydrophobic α-helix and a globular C-terminal domain exhibiting an immunoglobulin like fold. The N-terminal α-helices bundle to form the core of the filament, while the C-terminal globular domains face outside^12, 14–16^. The archaellum is not always homopolymeric, as a recent structure of *Methanococcus villosus* revealed a heteropolymeric assembly of the filament ^16^. For adhesion, *S. acidocaldarius* assembles the archaeal adhesion pili (Aap), which also belong to the T4FF ^17^. A comparison of known homologous structures of Aap from closely related thermoacidophilic organisms such as *S. islandicus* or *S. solfataricus* shows that these filaments have a high degree of glycosylation as a protective adaption to the highly acidic environment^18, 19^. In addition, Aap pili are also recognized by archaeal viruses as a potential virus receptor^18–21^. Another well-studied pilus is the UV-induced pilus, which is formed by various *Sulfolobus* species in response to DNA damage by e.g. UV light and leads to species specific cell-cell aggregation^22–24^. This allows the cells to exchange chromosomal DNA for homologous recombination ^25, 26^.

When Aap, UV pili and the archaella are deleted in *S. acidocaldarius*, the cells still form surface structures named “threads”. These are thin, 5 nm wide filaments that are highly abundant on the cell surface ^27^. Here, we present a 3.45 Å resolution structure of this filament and identify its so-far unknown subunit protein. The subunits are tightly interlinked via donor strand complementation and N-terminal isopeptide bonds, akin bacterial pili that are assembled via the Chaperone-Usher pathway. We find that the threads are highly glycosylated and based on our cryoEM map, we were able to model five complete N-glycan trees per subunit into our structure. Structure and sequence-based bioinformatic analysis reveals a so-far unknown archaea-specific class of cell surface filament.

## Results

### Threads are SPIII-independently assembled cell surface structure in *S. acidocaldarius*

Threads were already observed in the early 2000s ^17^ in *S. acidocaldarius* cells, however, their structure and assembly mechanism has been elusive. A triple knockout mutant of *S. acidocaldarius (ΔupsEΔarlJΔaapF*) that is unable to assemble archaella, UV pili or Aap pili, still forms threads^28^. Since the prepilin signal peptidase PibD (or class III signal peptide peptidase SPIII) is crucial for the assembly and function of archaella, UV-pili and Aap pili, we asked whether it is also essential for the assembly of the threads. However, a *ΔpibD* deletion mutant still showed thread filaments on its surface as analyzed by TEM (Supplementary Figure 1). Hence, threads are formed by a PibD independent pathway and thus do not belong to the T4FF superfamily.

### CryoEM and helical reconstruction of thread filaments

We sought to elucidate the structure of the thread filament by CryoEM. To this end, *S.acidocaldarius* strains either lacking archaella and Ups pili (MW158) or lacking PibD (MW114) were grown to stationary phase. The filaments were sheared from the cells and further purified by CsCl gradient centrifugation. The resulting sample containing a mixed population of threads and Aap pili was plunge frozen on cryoEM grids, of which 6272 cryoEM movies were recorded. Using CryoSPARC ^29^ contrast transfer function (CTF) correction and motion correction was performed, before filaments were automatically picked using the filament tracer program. The dataset was cleaned up by several rounds of 2D classification to remove erroneously autopicked Aap pili filaments and poorly resolved classes of threads (Supplementary Figures 2 and 3). From our 2D classifications it became apparent that threads often align into cables consisting of several parallel strands of thread filaments (Supplementary Figure 4). Similar cables were observed in micrographs of negatively stained *S. acidocaldarius* cells (Supplementary Figure 1), indicating that the threads are adhesive to each other and may enable adjacent cells to interact.

The Helix Refine routine within cryoSPARC allowed for low resolution 3D reconstructions of filaments without the need to impose helical parameters^29^. The resulting map with a resolution of 5.54 Å clearly showed that the thread filament consists of a helical string of subunits (Supplementary Figure 5) and allowed us to roughly determine the helical parameters in Real Space. A symmetry search around these values in CryoSPARC resulted in a clear peak at a rise of 31.6 Å, which was corroborated by a meridional reflection in Fourier power spectra calculated with SPRING, and a twist of –103.2° (Supplementary Figure 6) ^30^.

Applying these values to the 3D refinement resulted in a map of 4.02 Å resolution, which further improved to 3.45 Å after several rounds of CTF refinement. A local resolution estimation showed that the map was best resolved in the core of the filament where it reached 3.0 Å resolution (Supplementary Figure 7). The final map revealed that the thread filament consists of globular protein subunits that are arranged in a head-to-tail manner (Figure 1 A,D,E).

**Figure 1.**
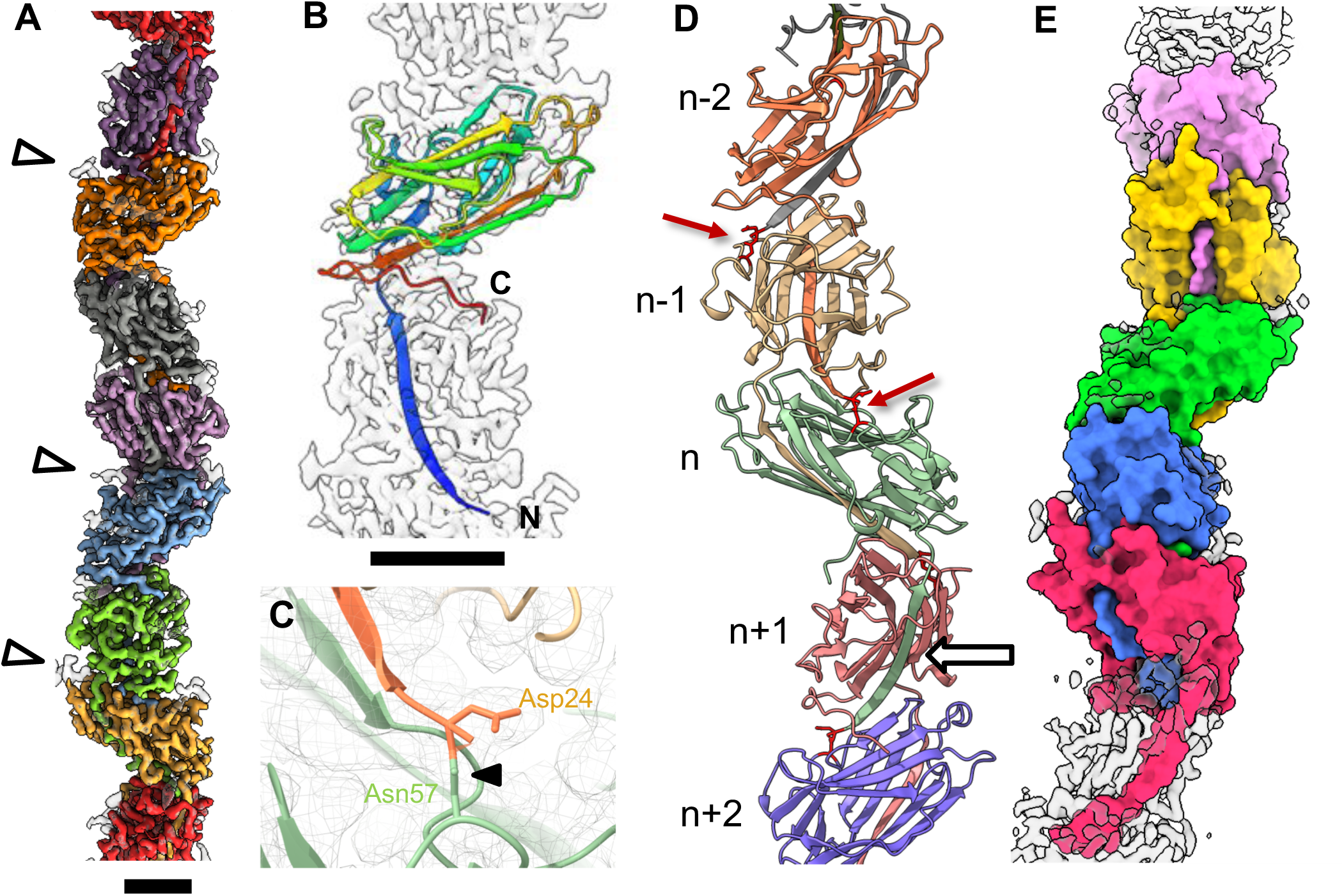
Helical reconstruction and atomic model of the Thread. **A,** Segmented surface representation of the cryoEM map showing each subunit in a different colour. Glycan densities are shown in white (white arrowheads). **B,** the atomic model of one Saci_0406 subunit (rainbow) fitted into the cryoEM map (transparent grey) N-terminus, blue; C-terminus, red. Glycans not shown for simplicity. **C,** intermolecular isopeptide bond. The N-terminal residue Asp24 of a subunit (n) is covalently bound to Asn57 two subunits along the chain (n+2). **D,** atomic model of 5 consecutive Saci_0406 subunits in ribbon representation. Glycans are not shown for simplicity. A white arrow indicates donor strand complementation between the N-terminal tail domain of subunit (n) and theC- terminal head domain of the next subunit along the chain (n+1). Red arrows indicate isopeptide bonds, highlighted in red. **E**, surface representation of five consecutive Saci_0406 subunits. The tail domain of each subunit n is partially buried in the two consecutive subunits n+1 and n+2. Scale bars, 20 Å

### Identifying the thread subunit protein

A cryoEM map at a resolution of 3.45 Å usually allows *ab initio* atomic model building if the protein sequence is known. We sought to determine the identity of the subunit via mass spectrometry, but the filament was resistant to digestion with trypsin or treatment with guanidine hydrochloride. This remarkable stability was also observed for the LAL/Aap pilus of *S. islandicus*, as well as for *S. solfataricus* which could not be denatured either by incubation with GdnHCl or boiling in HCL^18, 19^ . Hence, the subunit of the thread was identified from its structure, similar to previously described strategies (Supplementary Figure 8) ^19, 31, 32^.

We initially built a poly-alanine chain into our map to determine the direction and length of the backbone (Supplementary Figure 9). This revealed that the polypeptide sequence is approximately 206 amino acids long. We were able to pinpoint the position of characteristic amino acids along the backbone due to their distinctive shape and size. These included aromatic amino acids, as well as prolines. In addition, we recognised five glycosylation sites, which typically appear as distinct densities that protrude from the backbone and are too large to correspond to amino acid side chains. In *S. acidocaldarius,* N-linked glycosylation has been investigated biochemically and via mass spectrometry^40^. The glycans occur at conserved consensus sites with the sequence NXT/S and usually consist of a tri-branched hexasaccharide containing the unusual sugar 6-sulfoquinovose^40^. Using the position of these sequons, as well as those of our modelled characteristic amino acids as fingerprint, we blasted against the proteome of *S. acidocaldarius* for candidate proteins. This resulted in a single hit, Saci_0406 (Supplementary Figure 10).

Using the sequence of Saci_0406, we were able to build an atomic model for the thread unambiguously (Figure 1B-E). To corroborate our assignment of Saci_0406 as the filament subunit, we modelled the structure of this protein using Alphafold2. The resulting prediction closely matched our *ab initio* structure, (Supplementary Figure 11), indicating that Saci_0406 indeed forms the subunit of the thread filaments. Upon comparison of the predicted structure with our *ab initio* model of Saci_0406, we noticed that the latter lacked an N-terminal 𝜶-helix (Met1 – Ala23, Supplementary Figure 11) meaning that Saci_0406 is N-terminally processed. This was confirmed by analysing the sequence with the SignalP5 server which predicted a signal peptidase I processing site, cleaving after the N-terminal alpha helix between the position Ala23-Asp24 (AA sequence: …LVA/DV…) (Supplementary Figure 12) ^33^.

### Structure of the thread

Our atomic model revealed that each subunit consists of an N-terminal β-strand (tail) followed by a globular (head) domain. The head domain contains two β-sheets composed of 11 β-strands (Figure 1 B, D, E). On the subunit level, one β-strand is missing to complete the sheet (Figure 1, Supplementary Figure 13), leaving a gap between β-4 and β-12. Upon closer investigation of the assembled filament, this missing strand is replaced by the β1 tail strand of the previous subunit along the filament (n-1) (Supplementary Figure 13B). Each head domain hereby forms a mainly antiparallel beta-blade of a mixed type, comprising 12 beta-strands in total (Figure 1D, Supplementary Figure 13). Such donor strand complementation has previously been observed in bacterial T1P, P-pili and T5P, where it has been proposed to contribute to the stability of the filament (Figure 4)^34, 35^. In the thread, the β-tail extends further into the subunit two positions along the filament (n+1; Figure 1D, Supplementary Figure 13 B). β-1 of the subunit n is partially buried in the subunits n+1 and n+2 (Figure 1E), further adding to the stability of the interaction. Moreover, Asn57 of each subunit forms an unusual isopeptide bond with the N-terminus (Asp24) belonging to a protomer two positions along the filament (n+2) (Figure 1C, D).

### The thread is highly glycosylated

Along the map of the *S. acidocaldarius* thread, we found dead-end protrusions that did not resemble protein backbone nor side chains. These protrusions were exclusively found at asparagine residues, N56, N80, N83, N121 and N146 of each thread subunit (Figure 2). Each of these asparagine residues are part of an N-glycosylation consensus sequon (N X S/T). Since it is well known that surface exposed proteins are highly glycosylated in *S. acidocaldarius*, we concluded that these densities correspond to N-glycan trees ^36^. The glycan density associated with Asn146 was particularly well resolved, presumably due to its unusual parallel position to the backbone, which likely reduces its flexibility (Figure 2F). Based on these densities, we were able to model the complete sequence of previously confirmed tribranched hexasaccharide moiety of *S. acidocaldarius* into the filamentous structure (Figure 2) ^37^. Connected to each of the five glycosylation sites at the corresponding asparagine, are two N-acetylglucosamine (GlcNAc) residues, the second of which branches to bind two individual Mannose (Man), Glucose and one 6-sulfoquinovose molecule.

**Figure 2.**
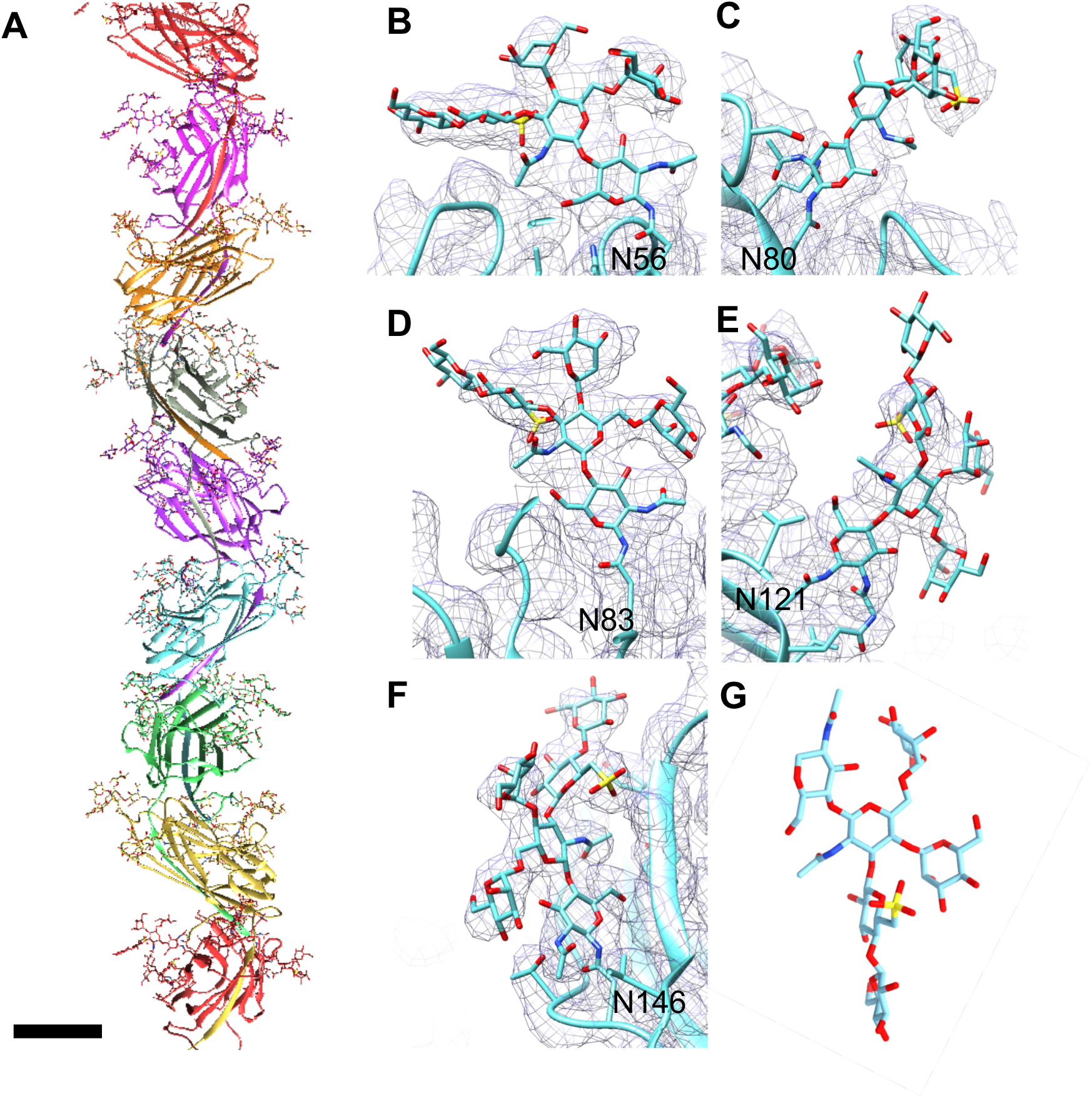
Threads are highly glycosylated. **A,** atomic model of the thread filament. Backbone in ribbon and glycans in stick representation. **B - F,** closeups of the five glycosylation sites found in Saci_0406. Atomic models of the glycans are in stick, the polypeptide backbone in ribbon representation. The CryoEM density map is shown as grey mesh. **G,** structure of the *S. acidocaldarisus* glycan in stick representation. Scale bar in A, 20 Å

While glycosylation sites were previously only partially resolved observed in structures of crenarchaeal pili^18, 19^, our structure allowed us to model the full N-glycan tree of *S. acidocaldarius* throughout the entire filament (Figure 2A).

### Threads are conserved among some Crenarchaea

Asking how widespread these threads are in archaea, we blasted Saci_0406 and found candidate proteins with a high sequence similarity in various other crenarchaea. A structural homology search using the NCBI protein blast server revealed that proteins with a similar predicted structure to that of Saci_0406 are encoded in the genomes of various *Vulcanisaeta* and *Acidianus* species (Figure 3). Some homologs are forming a cluster of orthologs (arCOG10215) comprising members from the crenarchaeal phyla^38^. Predicting their structures using Alphafold2 revealed that these homologues are almost structurally identical to Saci_0406 (with average RMSDs of 2-3 Å; Figure 3 B-E). All five homologues follow the same beta blade head structure as the thread subunit Saci_0406. The first 7 amino acids after after the N-terminal processing by the signal peptidase 1 are also identical, including Asp24 and the YYY motif (Supplementary Figure 14). Based on the conserved Asp24, it is intriguing to hypothesize that the isopeptide bond is conserved in the thread homologues.

**Figure 3.**
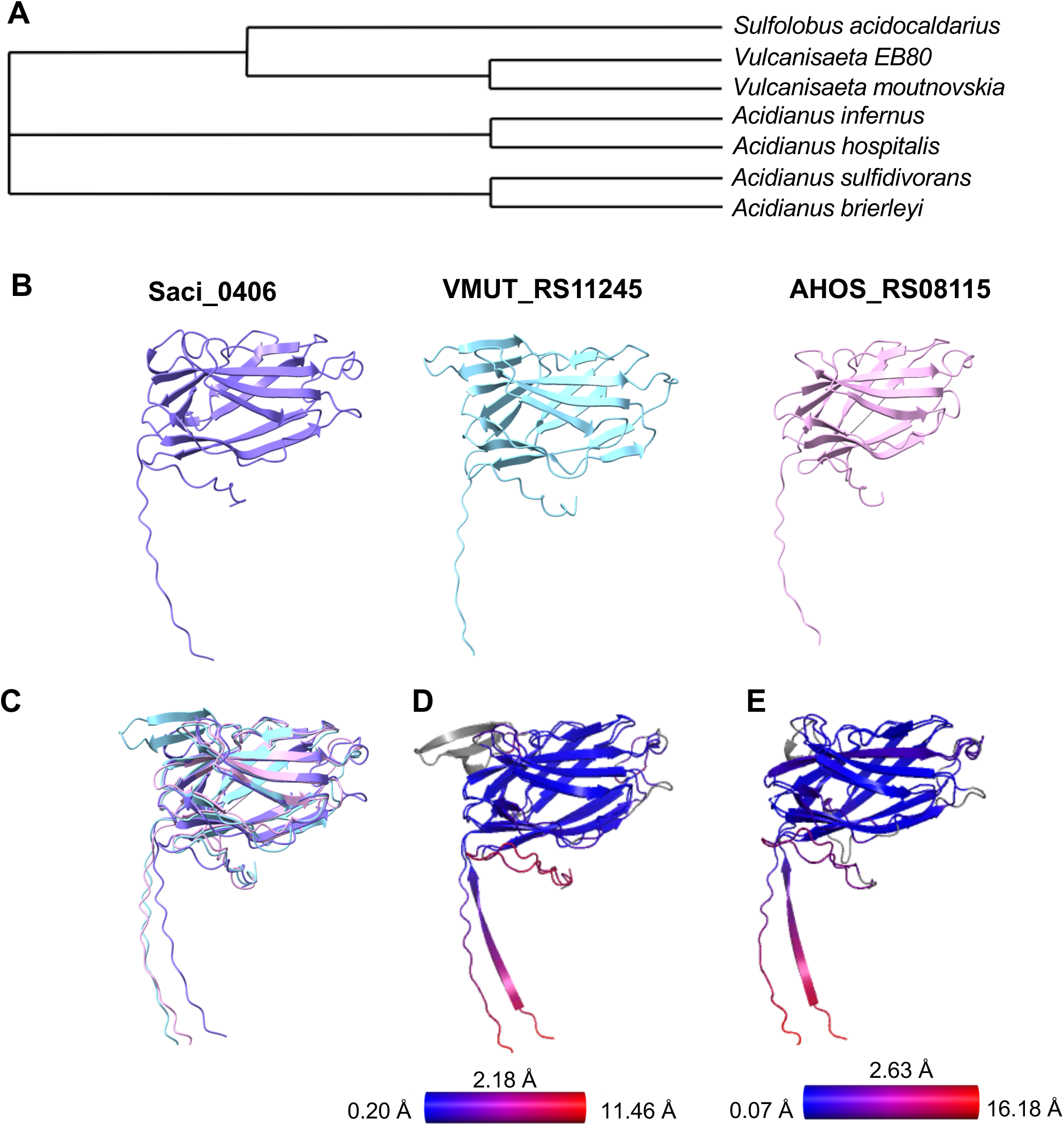
Threads are conserved in crenarchaeal relatives. **A,** Phylogenetic tree of species where *Saci_0406* homologs have been identified. **B,** Saci_0406 (purple) compared with Alphafold2 predictions of homologs found in *V.moutnovskia* (blue) *A.hospitalis* (pink) **C,** superimposing the structures from B shows their similarity. **D, E,** pairwise RMSD values calculated between Saci_0406 and the thread homologues from *V.moutnovskia* (**D**) and *A.hospitalis* (**E**). Blue indicates low and red high values. Average value RMSD is 2.18 Å for D and 2.63 Å for E.

Following on from this, we analysed the syntheny of *saci_0406*. Orthologs of *saci_0406* co-occur with those of *saci_0405*, *0406, 0407 and saci_0408,* which has its own promoter (Supplementary Figures 15, 16). Indeed, the genetic neighbourhood in closely related species seems to be very conserved. In some strains, the genes orthologous to *saci_0407* and *saci_0408* are fused in one gene, whereas in others it seems that one is a noncoding region (Supplementary Figure 16).

### A putative cap protein

As blasting the corresponding protein sequences did not yield any conclusive results, we predicted the structures of Saci_0404, 0405, 0407 and 0408 using Alphafold2 (Supplementary Figure 17). While the predictions for Saci_0404, 0407 and 0408 did not lead to conclusive insights, the one for Saci_0405 gives rise to a hypothetical role in thread assembly.

The predicted structure of Saci_0405 has a remarkably similar fold compared to Saci_0406 (Supplementary Figure 18 A), including the extended tail. The mean RMSD between the structure of Saci_0406 and the Alphafold prediction of Saci_0405 measured only 4.58 Å (Supplementary Figure 18 B) and an Alphafold2 prediction of both proteins in a complex placed the N-terminal tail of Saci_0405 into the donor strand acceptor site of Saci_0406 (Supplementary Figure 18 C). Modelling Saci_0405 into a terminal position of our thread structure revealed that Saci_0405 could complement the β1 acceptor site of a Saci_0406 n+1 subunit (Supplementary Figure 18 F,G). However, compared to the experimental structure and the Alphafold-2 prediction of Saci_0406, Saci_0405 lacks a donor strand acceptor site in its globular domain (Supplementary Figure 18 D). This suggests that Saci_0405 could function as a filament cap. In line with this hypothesis, Saci_0405 and Saci_0406 are very likely expressed under one promoter (Supplementary Figure 19), indicating co-expression and thus a functional link between the two proteins. Moreover, the RNAseq data show that *saci_0405* is expressed at lower levels and thus in smaller copy numbers than *saci_0406*, in accordance with a putative function as a cap ^39^. Cap proteins are a hallmark for bacterial filaments that assemble through donor strand complementation. All these filaments require a terminal subunit that shields the terminal tail domain and itself is not dependent on a downstream protein to provide a completing 𝛃-strand^2^.

### Threads resemble bacterial Type I pili assembled by the Chaperone Usher pathway

The overall architecture of the *S.acidocaldarius* thread filament is reminiscent of bacterial bacterial Saf pili and the recently characterized *P. gingivalis* type-V pili ^34, 35, 40^ (Figure 4 C-E). Threads, Saf and T5P all consist of stacked subunits that are interlinked by donor strand complementation of the N-terminal β-strand. However, on the subunit level, there is little structural similarity between Threads and bacterial Saf and T5P pili.

**Figure 4.**
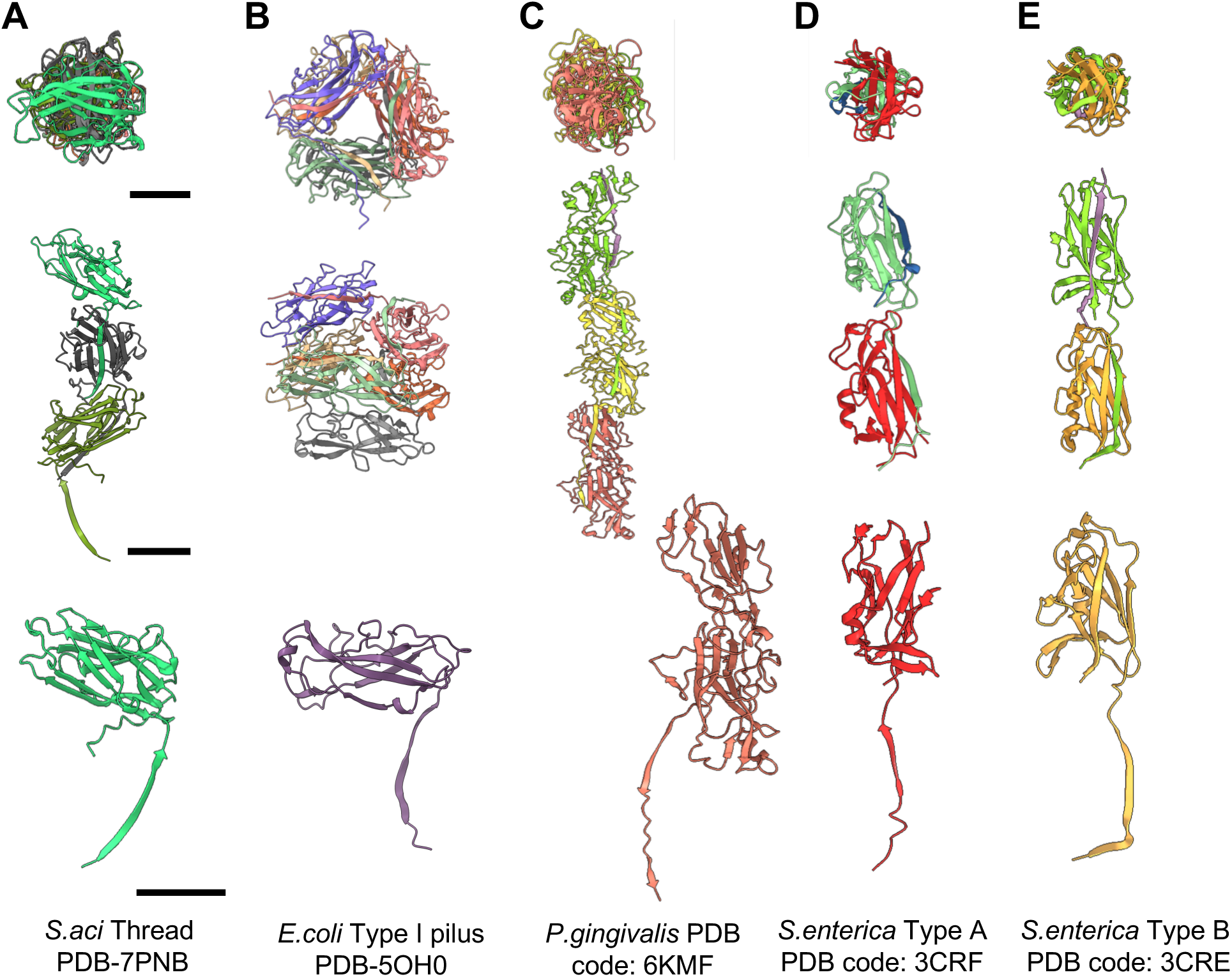
Comparison between threads and bacterial pili with donor strand complementation. **A-E,** The structures of the *S. acidocaldarius* Thread (A), *E.coli* T1P (B), *P. gingivalis* T5P (C) and *S. enterica* Type A (D) and Type B Saf pili (E). Top panel, filament cross section; middle panel, filament side views. Bottom panel, subunit structure for each filament. **A**, *S. acidocaldarius* thread (Saci_0406; PDB-7PNB); **B**, *E.coli UPEC J96* Type I pilus (PDB- 5OH0); **C.** P. gingivalis Type V pilus (FimA1;PDB-6KMF). D. S. enterica Typhimurium Saf Pilus type A (PDB:3CRF). E. *S. enterica* sv. Typhimurium Saf Pilus type B (PDB:3CRE). Scale bars, 20 Å

When comparing threads and bacterial T1P, there is a significant structural difference in the filaments’ general architecture (Figure 4 A,B). Whereas threads can be described as linear 40 Å wide, beads-on-a-string like filaments, T1P and P pili form hollow (likely solvent-filled) tubes with diameters of up to 60 Å ^41, 42^. This is reflected by distinct helical parameters. Whereas the threads have a rise of 31.6 Å and twist of –103.2°, Type I pili have a reported rise of 7.7 Å and a twist of 115°, type P pili show close similarity with values of 7.7 Å and 109.8° for rise and twist respectively ^41, 42^. The type V pilus has a rise of 66.7 Å with a twist of 71° ^34^.

However, comparing the structure of the thread subunit Saci_0406 with those of bacterial pilins reveals a striking similarity with bacterial T1P and P pilins of UPEC and UTI strains of *E.coli* (Figure 4 A, B). Superimposing the structure of Saci_0406 with those of FimA and PapA of *E. coli* yields overall RMSD values of 9.94 – 11.76 Å with a structurally highly conserved β-blade core (RMSD values around 2 Å; Supplementary Figure 20). Notably, T1P and P pili assemble via donor strand complementation, catalyzed by the chaperone-usher pathway ^41–45^. However, despite the structural similarity between Saci_0406 and the bacterial Type I pilins, multiple sequence alignments indicated low sequence homology (Supplementary Figure 21). Thus, we hypothesize that donor strand complementation and ß-sheet rich globular domain is a convergently evolved structural solution to build a remarkably stable filament in the harsh thermoacidic environment that *S. acidocaldarius* inhabits.

## Discussion

Extracellular filaments are often involved in adhesive or propulsive processes and thus have to be resilient to mechanical forces and adverse or changing environmental conditions. This is especially true for the threads of *S. acidocaldarius*, which inhabits environments with highly acidic pH and temperatures not far from the boiling point of water. Threads likely gain particular tensile strength and resilience against heat and acidic pH through three structural adaptions: Donor strand complementation, intermolecular covalent isopeptide bonds and a high degree of glycosylation.

Donor strand complementation has previously been found in various types of bacterial pili, such as T1P, P-pili and T5P^2^, as well as in the very recently published structure of the archaeal bundling pili of *Pyrobaculum calidifontis* ^46^. All of these filaments have been implicated in adhesion or biofilm formation. In accordance with this, adhesion experiments with *S. acidocaldarius* mutant that lacked archaella and Aap but still expressed threads retained 25 % adherance compared to WT. In addition, this mutant strain was still able to form biofilms and even showed a higher density of biofilm-borne cells compared to WT ^28^. We show that threads tend to align into cables of several of filaments, suggesting a propensity to connect neighbouring cells. It is intriguing to hypothesize that threads are necessary for scaffolding and spatial organization of the biofilm.

Threads appear to be widespread in the archaeal domain, as demonstrated by our finding of *saci_0406* homologs across various crenarchaea (ArCOG10215). Multiple sequence alignments revealed a conserved SP I cleavage site in all homologs (Supplementary Figure 14). Moreover, Alphafold2 predictions of homologs from *Acidianus hospitalis* and *Vulcanisaeta moutnovskia* reveal a very similar fold (Figure 3). The N-terminal ß-strand that is involved in donor strand complementation is similarly conserved, indicating that it is a shared trait in crenarchaeal threads.

Threads are covalently interlinked by a non-canonical isopeptide bond between Asn57 of subunit n and the N-terminal Asp24 of the subunit n+2. Strikingly, Asp24 is conserved throughout all homologs identified (Supplementary Figure 14) . Similar isopeptide bonds have been shown to provide additional stability in filaments of some Gram-positive bacteria, but these usually occur between Lys and Asn/Asp residues ^47–49^.

In sortase mediated pili of Gram-positive species *Actinomyces oris*, an isopeptide linkage between Thr499 and Lys182 ^50^ is found and in *Streptococcus pyogenes,* the C-terminal Thr311 is covalently bound to Lys161^51^. In both cases, the isopeptide is established between directly adjacent (n and n+1) subunits, in contrast to the isopeptide bond in the thread that established between the subunits n and n+2.

In bacteria, the formation of the isopeptide is catalysed by a sortase enzyme, which recognises a specific motif near the C-terminus on newly incorporated subunits and cleaves the terminus that establishes the isopeptide with a upstream subunit^50^. A well-studied archaeal sortase is the archaosortase ArtA from the euryarchaeon *Haloferax volcanii,* which is essential for anchoring cell surface proteins to the cell membrane. Despite similar catalytic residues, no further homology is found between ArtA and bacterial sortase enzymes^52^. As sortases have so far not been found in crenarchaea, it is likely that either a yet to be identified transpeptidase aids the formation of the isopeptide bonds, or that they form spontaneously. Indeed, spontaneous isopeptide bond formation has been found to occur in Gram positive bacteria ^53^.

As no isopeptide bonds were reported in the recently solved archaeal bundle pili of *P. calidifonitis*, they may be a unique feature of threads^18, 19, 46^. In addition, compared to the archaeal bundle pili, Saci_0406 does not contain a PFAM domain, which appears to be confined to Caldarchaeales and an uncharacterized lineage^46^. These differences suggest that threads form a separate class of archaeal cell surface filaments.

Each thread subunit is glycosylated at five asparagine residues. Based on our cryoEM map, we were able to build the complete structure of the N-glycan of *S. acidcaldarius*, which was previously sequenced via mass spectrometry ^37^. In Archaea, glycosylation is highly diverse, with a large a variety of possible sugar moieties that may have additional modifications^3^. A high degree of glycosylation is a common feature of surface exposed archaeal proteins. For *S.acidocaldarius*, glycosylation has been shown to be crucial for motility, proper S-layer function, as well as species-specific recognition^22, 54^. In addition, a high degree of glycosylation is likely an important adaptation to highly acidic environments, and contributes to the remarkable stability of archaeal filaments, as shown for the Aap of *S. solfataricus* and *S. islandicus* ^18, 19^.

The structure of the thread suggests key steps in its biogenesis, as shown in Figure 5. Bioinformatics indicates that the thread subunit Saci_0406 is secreted via the SEC-pathway. In the next step, the archaeal signal peptidase I will recognize the signal peptide and cleave the hydrophobic, membrane bound N-terminus of the pre-protein. This will release the mature Saci_0406 protein with an N-terminal Aspartate and an archaea-specific triple Y motif after cleavage site ^55^ as a soluble protein into the pseudo-periplasm. Saci_0406 will then be glycosylated by AglB, the oligosacharyltransferase which transfers the N-glycan to the specific aspartic acids within the N-glycsylation sequon^56^.

**Figure 5.**
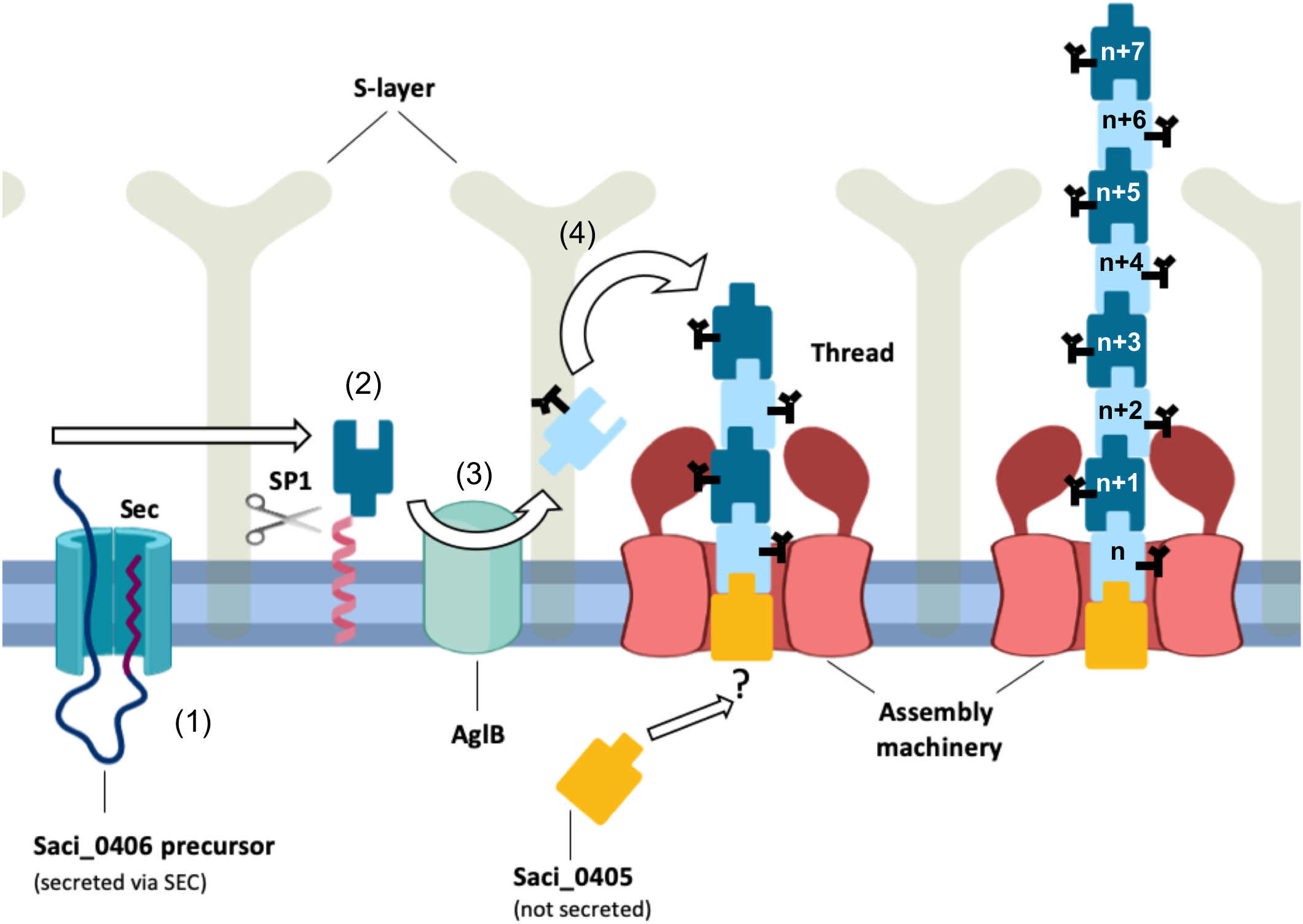
Hypothetical models for thread assembly. *S. acidocaldarius* precursor proteins are initially secreted as membrane proteins into the cell membrane (1). Signal Peptidase 1 (SP1) cleaves the N-terminal signal peptide, including the transmembrane helix (2). The protein is glycosylated by AglB (3; glycans shown as black sticks). Soluble, mature Saci_0406 proteins assemble into the thread via donor strand complementation (4). This process may be spontaneous or catalyzed by an assembly machinery. The putative cap protein Saci_0405 is not secreted through SEC and thus likely remains in the cytoplasm. How it interacts with the cell-proximal end of the thread is unknown.

The presence of donor strand complementation and the structural similarity with bacterial T1P poses the intriguing question if the thread filament is assembled via a process akin the bacterial Chaperone Usher pathway. Here, pre-pilins are initially bound by the chaperone (FimC in Type I and PapD in P pili of *E. coli*) at the nascent N-terminus. The donor strand of the chaperone stabilizes the pilin subunit until it is integrated into the growing pilus filament at the usher pore ^2, 57–59^.

In contrast, in T5P, precursors of the subunit proteins are lipidated at a cysteine in a lipobox region. The precursors are then guided to the secondary membrane, likely by the Lol pathway, and after subsequent cleavage by an arginine/lysine specific protease the filament is polymerized^60^. In the recently published structure of the Gram positive *B. subtilis* TasA filament, the TasA subunit tightly interact via donor strand complementation forming a thin filament. In the monomeric form TasA stabilizes itself by self-complementing the binding site for the donor strand. In the filament, TasA undergoes a conformational change and the binding site is free for donor strand complementation from the adjacent subunit. The filament polymerization is initialized by an accessory protein TapA which can bind to the strand accepting groove of TasA, so TasA can initiate binding to the next subunit ^61^. Structural prediction of the archaeal bundle pili subunit revealed a similar mechanism ^46^. In contrast, we did not observe auto complementation for Saci_0406, potentially highlighting a distinct assembly mechanism.

Analysing the gene cluster surrounding Saci_0406, did not reveal proteins that could definitively be assigned to either a chaperone or an usher protein. In line with this, *S. acidocaldarius,* and in fact most known archaea, do not possess an outer membrane, suggesting that an usher protein may not be necessary. Instead, many archaea, including *S. acidocaldarius* are encased by a proteinaceous and highly porous cell wall, called S-layer ^17^. The S-layer of *S. acidocaldarius* contains triangular and hexagonal pores with reported diameters of ∼45 Å or ∼80 Å, respectively ^62^. With a diameter of 40 Å, threads would comfortably fit through the 80 Å hexagonal pore, which may thus serve as a conduit for the thread at the cell surface. In line with this, S-layers have previously been shown to be involved in the anchoring of archaella ^63^.

Saci_0405 has a similar fold to Saci_0406 with a N-terminal ß-strand but no donor strand acceptor site. This means that Saci_0405 would be able to provide its ß-strand for a downstream Saci_0406 molecule without needing a complementing ß-strand upstream. Hence, it is likely that Saci_0405 forms the initial nucleating subunit for the Saci_0406 thread. Similar observations were made for the TasA filament formed by *B. subtilis*, where TapA a helps nucleating filament assembly of TasA, as well as the recently characterized archaeal bundle pili ^46, 61^. Interestingly, we find that unlike Saci_0406, the putative cap protein Saci_0405 does not have a predicted SEC signal sequence or a transmembrane domain. This suggest that Saci_0405 likely remains in the cytoplasm and acts as putative capping protein that may anchor the threads to a yet to be identified assembly platform (Figure 5).

Taken together, we propose that archaeal threads comprise a new class of archaeal biofilm-forming protein filaments that have arisen independently from similar bacterial pili. We suggest that threads are assembled by yet unknown pathway, in which donor strand complementation provides the driving force for filament assembly. To shed more light on the assembly mechanism of the threads, the assembly machinery will have to be identified and characterised in future studies.

## Acknowledgements

We acknowledge Diamond for access and support of the cryoEM facilities at the UK national electron bio-imaging centre (eBIC), funded by the Wellcome Trust, MRC and BBSRC.

We would like to thank the EM facility at the Faculty of Biology at the University of Freiburg for access to their microscopes. The TEM (Hitachi HT7800) was funded by the DFG grant (project number 426849454) and is operated by the University of Freiburg, Faculty of Biology, as a partner unit within the Microscopy and Image Analysis Platform (MIAP) and the Life Imaging Center (LIC), Freiburg. We also thank Carsten Sachse at Forschungszentrum Jülich, Germany, for his helpful suggestions regarding helical reconstruction.

MG, MM and BD have received funding from the European Research Council (ERC) under the European Union’s Horizon 2020 research and innovation program (grant agreement No 803894). VG and AN have received funding from the UK’s Biotechnology and Biological Science Research Council (grant agreement number BB/R008639/1).

SS And SVA were supported by the Collaborative Research Centre SFB1381 funded by the Deutsche Forschungsgemeinschaft (DFG, German Research Foundation)—Project-ID 403222702—SFB 1381. SS and SVA were also funded by the Deutsche Forschungsgemeinschaft (DFG, German Research Foundation) under Germany’s Excellence Strategy (CIBSS – EXC-2189 – Project ID 390939984).

## Materials and Methods

### Cell growth and threads isolation

*S acidocaldarius* MW158 or MW114 were inoculated from cryostock in 6 x 5 mL basal Brock (pH3) supplemented with 0.1 % NZ-amine, 0.2 % dextrin and 10 µg/mL uracil. The cells were grown for two days at 75 °C with light agitation. 5 mL of preculture was inoculated per litre main culture. The cells were grown for 48 h until they reached an OD600 =1. Afterwards cells were harvested using 5000 x g for 25 min at 4°C. The cell pellets from 2 litre main culture were resuspended in 20 mL Basal Brock (pH 3) without FeCl_3_. To shear the threads either shearing was done as described in Henche et al 2012 ^27^ or a peristaltic pump (Gilson Minipuls) was used which was connected to syringe needle with 1.10 mm in diameter and 40-50 mm in length (Braun GmbH). The cells were homogenized at 25 rpm for 1h. The syringe needle were switched to more narrow ones with 0.45 mm in diameter and 10 mm and shearing was done at 25 rpm for 1h. Subsequently, the sheared sample was centrifuged at 12 000 x g for 25 min at 4°C. The supernatant was subjected to ultracentrifugation at 200 000 x g for 90 min, at 4°C. The resulting pellet was resuspended in 500 µL Basal Brock without FeCl_3_ and layered on 4.5 mL CsCl_2_ (0.5 g/mL) and subjected to density gradient centrifugation at 250 000 x g for 16 h at 4°C. A white band in the upper third of the tube was detected and collected. This was diluted to 5 mL in Basal Brock without FeCl_3_ and pelleted at 250 000 x g for 1h at 4°C. The pellet was resuspended in 150 µL of citrate buffer (25 mM sodium citrate/citric acid, 150 mM NaCl, pH 3) and stored at 4 °C.

### Negative stain transmission electron microscop

5µL of cells were applied on freshly glow-discharged 300 mesh carbon-coated copper grids(Plano GmbH, Wetzlar Germany) and incubated for 30 s. The excess liquid was blotted away and grids were stained with 2% uranyl acetate. Imaging was either performed with a Zeiss Leo 912 Omega (tungsten) (Carl Zeiss, Oberkochen, Germany) operated at 80kV, equipped with a Dual Speed 2K-On-Axis charged coupled device (CCD) camera TRS, Sharp-Eye (TRS Systems, Moorenweis, Germany) or Hitachi HT8600 operated at 100 kV equipped with EMSIS XAROSA CMOS camera, or a FEI T12 (Eindhoven, The Netherlands) operated at 120 kV equipped a LaB6 filament and a Gatan Oneview CMOS detector (Pleasanton, USA).

### Cryo-EM sample preparation and data collection

A 3 μl drop of suspension containing a mixture of Aap pili and threads was applied to glow-discharged 300 mesh copper R2/2 Quantifoil grids. The grids were blotted with 597 Whatman filter paper for 5 sec, using -1 blot force, in 95 % relative humidity, at 21 °C and plunge-frozen in liquid ethane using a Mark IV Vitrobot (FEI). Image data were collected using a FEI Titan Krios electron microscope, operating at 300 kV in nanoprobe mode using parallel illumination and coma-free alignment, on a Gatan K2 Summit electron detector in counting mode at a calibrated magnification of 134,048× (corresponding to a pixel size of 1.047 Å at the specimen level) with defocus range from −2.0 to −3.5 µm using 0.3 µm steps. The microscope and camera were controlled using EPU software (Thermo Fisher Scientific). Images were recorded as movies at a dose rate of 0.77 e/Å^2^s^−1^ at 40 frames s^−1^, 10 s exposure, with an accumulated total dose of 42.33 e/Å^2^. Cryo-EM statistics are presented in Supplementary Figure 13.

### Cryo-EM image processing

Frames from 6272 movies were aligned using full-frame motion correction as part of the cryoSPARC ^29^ package to correct for stage drift. Patch CTF estimation was then used to estimate defocus variation. The e2helixboxer program from EMAN2 ^64^ was used to manually pick filaments so that 2D classes could be quickly generated in Relion ^65^ . From here the 2D classes were imported into cryoSPARC ^29^, where the Filament tracer program picked out 2,243,141 filament fragments comprising of both AAP filaments as well as Threads. Particle coordinates were extracted from the micrographs with a box size of 512 × 512 pixel covering 10-11 Thread subunits within a circular mask. 5 rounds of 2D classification utilising 50 classes each separated the Aap pili from the threads leading to a total of 188,620 particles from the best classes. Helix Refine was used to create a non-biased 3D volume which could then be examined to determine the helical parameters. Using Chimera ^66^ to visualise the filament at a resolution of 7.79 Å, a rise of approximately 90 Å was predicted in real space. This was then confirmed using the symmetry search utility in cryoSPARC ^29^ specifying a rise range of 80-120 Å and a twist range of –180° to +180°. A sharp peak was found with a rise of ∼95 Å and a twist value of ∼40° which was used to refine the non-biased helix refine job, leading to a map with resolution of 4.30 Å. Closer observation of this volume revealed that the above-mentioned parameters corresponded to not 1, but 3 repeating subunits. lllSymmetry search was then carried out again in cryoSPARC ^29^, using this 4.30 Å map, to reveal the final parameters of –103° and 31.6 Å for twist and rise respectively. From here, the iterative process of performing a Local motion correction, Local CTF Refinement and Global CTF Refinement job followed by a Helix Refine job was performed, until resolution of our map reached 3.46 Å. The resolution was estimated using a Fourier shell correlation between two independently refined half sets of data using a Fourier shell correlation of 0.143. This final map was then denoised and postprocessed using DeepEMhancer ^67^ . The local resolution calculated with Local Resolution Estimation in cryoSPARC ^29^, and mapped in ChimeraX ^68^ (Supplementary Figure 7).

### Model building and validation

A polyalanine model was built into the observed density using Coot ^69^ . Side chain identities were assigned for uniquely shaped aromatic, proline and low-density glycine residues. Glycosylation sites in *Sulfolobus acidocaldarius* are predicted to be N-glycosylation and thus such sites were labelled as Asn x Ser/Thr motifs. ‘X’ residues were modelled as alanines or serines. Medium sized residues pointing inside the beta barrel head structure, were modelled as aliphatic leucine, isoleucine, or valine. Short surface residues were modelled as asparagine, serine or aspartate. The resulting putative amino acid sequence was run through PSI-BLAST search ^70^ against the *S. acidocaladarius* genome on the NCBI BLAST server ^71^ . The only protein which matched the search pattern with high E-value of 5e-17 (28 % sequence identity over 62% of length) was Saci_0406. The sequence of Saci_0406 could subsequently be unambiguously modelled into our map. An unusual bridging feature proved to be an isopeptide link formed between the main chain nitrogen of the N-terminal Asp24, as predicted by SignalP-5.0 server^33^ and the side chain of Asn57 from the N-2 chain in the filament. To confirm our model, we predicted the structure of Saci_0406 in AlphaFold2 ^72^, which resulted in an almost precise match for the head domain (Supplementary Figure 10). The remaining monomers in the filament were positioned in density by phased molecular replacement implemented in MOLREP^73^ adapted for EM. Later CCP4^74^ script was prepared to propagate changes in the rebuilt monomer to others in the filament. The glycan structures were modelled using Coot^69^, dictionaries for isopeptide link and unusual sugars were prepared using JLIGAND^75^ . The structure was refined using REFMAC5^76^ in the ccpem interface.

### Structural prediction (Alphafold2)

The genome of *S. acidocaldarius* was visualised using several tools. The KEGG genome database^77^ allowed us to accurately determine which proteins surrounding Saci_0406 were likely to be part of the same operon. Syntax^78^ was used to search for the Saci_0406 sequence to see the corresponding genomes in other archaeal species, to search for homologous gene clusters. From these sources, we concluded that Saci_0404 – Saci_0409 were likely to within the same gene cluster. Alphafold2^72^ was used to predict the structures of Saci proteins 0404-0409, using the online ColabFold^79^ tool. Results were observed in Chimera^66^ and ChimeraX^68^ to determine if any relationships between the different protein members could be determined. PDBefold^80^ as well as predictions of other archaeal proteins found using Syntax were observed as stated above. Sequence analysis was done using Clustal Omega^81^ to observe sequence similarities.

### 3D variability analysis and molecular flexibility

3D variability was conducted in cryoSPARC^29^ to determine the molecular flexibility of the Threads. The corresponding image files generated were used to create movies made up of 20 frames each visualized USFC Chimera^66^ . The refined Saci_0406 model was positioned in the frame 0 map using MOLREP^73^ . Subsequently, this model was refined by real space refinement implemented in Phenix^82^ against the maps corresponding to the first or last frame of the movie.

### Sequence Analysis

Two iterative PSI-BLAST^70^ searches were done using the sequence of Saci_0406 to determine over the largest sequence cover, the largest percentage identity for foreign species. The top five differing results were then structurally compared against each other using Clustal Omega^81^ .

### Structure analysis and presentation

Already published filaments were assessed for structural similarity to the threads. *P. gingivalis* FimA1 and *S. enterica* Saf Pilus both show beta sheet inclusion between adjacent subunits in a conceptionally similar method to the threads and so comparisons were made using USFC Chimera^66^ of both the filament and monomeric forms. Several *E. coli* Type I pili were also studied to highlight the similarities monomerically to the threads.

### Data deposition

The atomic coordinates and electron density map were deposited in the Protein Data Bank (https://www.rcsb.org/) with accession number 7PNB and in the EM DataResource (https://www.emdataresource.org/) with the accession number EMDB–13546.

## Competing interests

The authors declare no competing interests

---

**Supplementary Figure 1.**
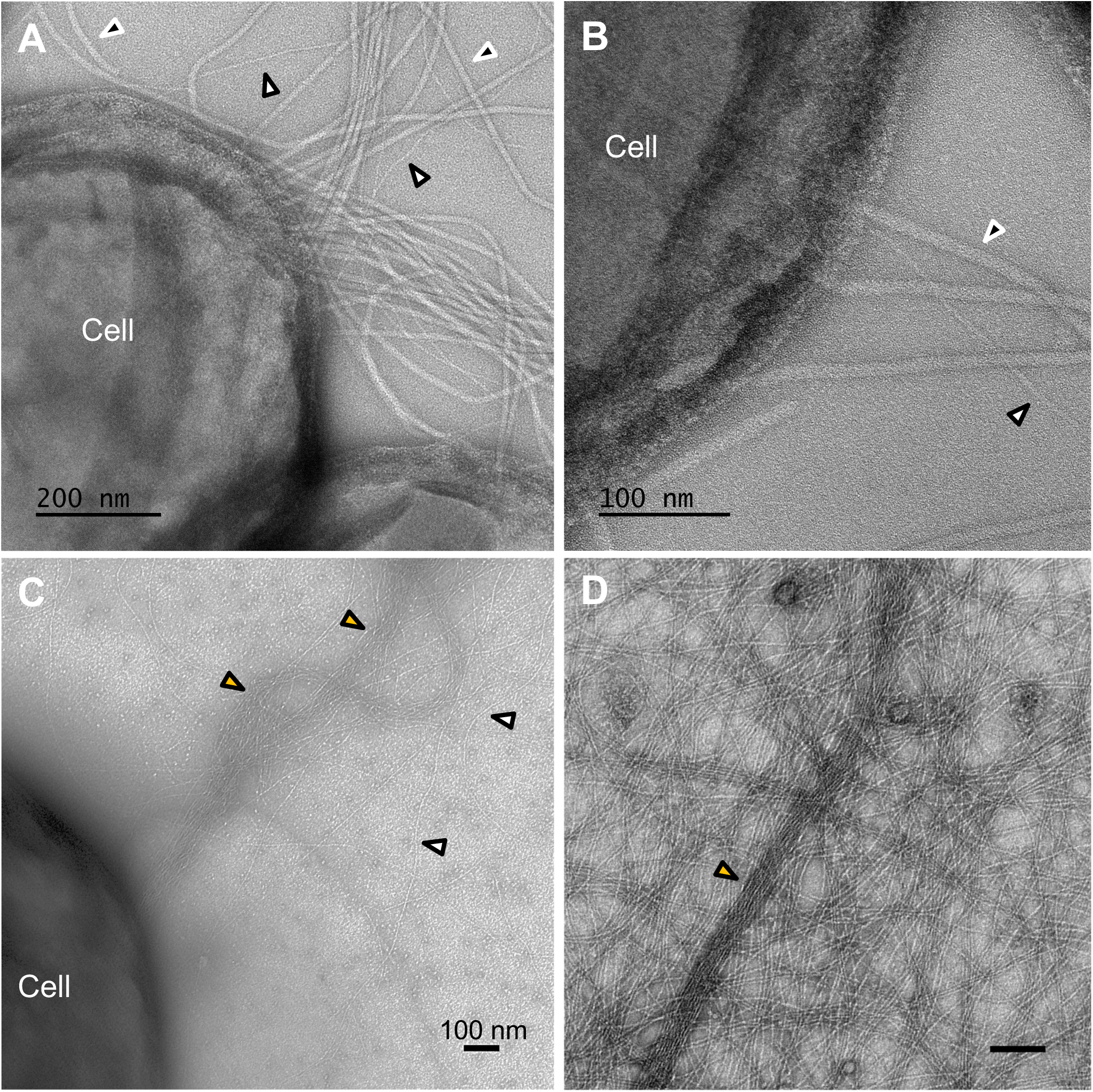
Negative Stain transmission electron microscopy *S. acidocaldarius* MW001 and *ΔpibD* mutant. **A**, **B**, micrographs of the cellular periphery of *S. acidocaldarius* MW001 cells at different magnifications. White arrowheads, Threads; black arrowheads Aap or archaella. **C**, micrograph of *ΔpibD* mutant that lacks archaella and Aap but retains threads. Orange arrowhead indicates a cable of threads. **D**, micrograph of isolated Thread filaments individual filaments and cables are evident in the purified filaments.

**Supplementary Figure 2.**
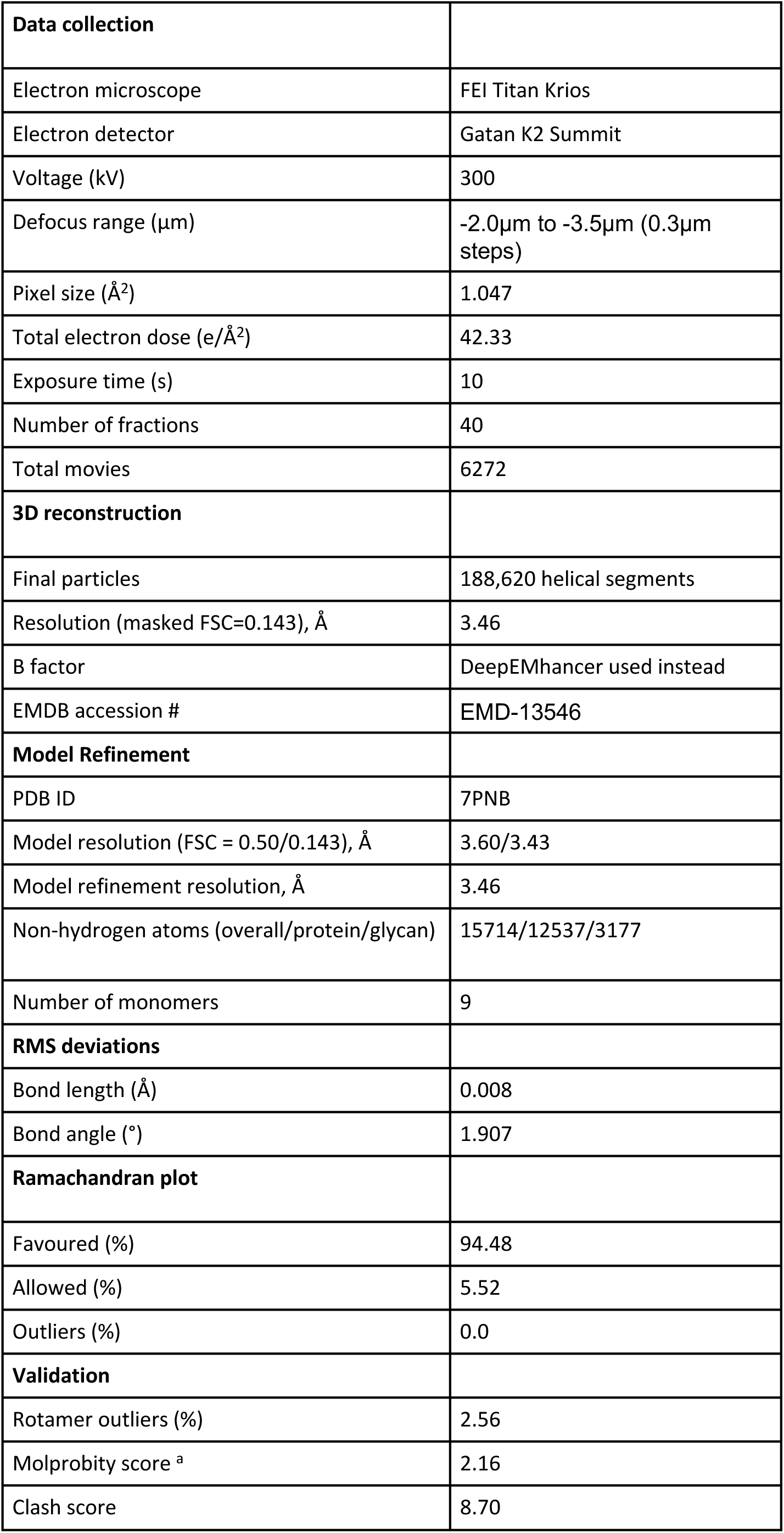
CryoEM and model building statistics. Table illustrating details on microscopy, data collection, image processing and model validation.

**Supplementary Figure 3.**
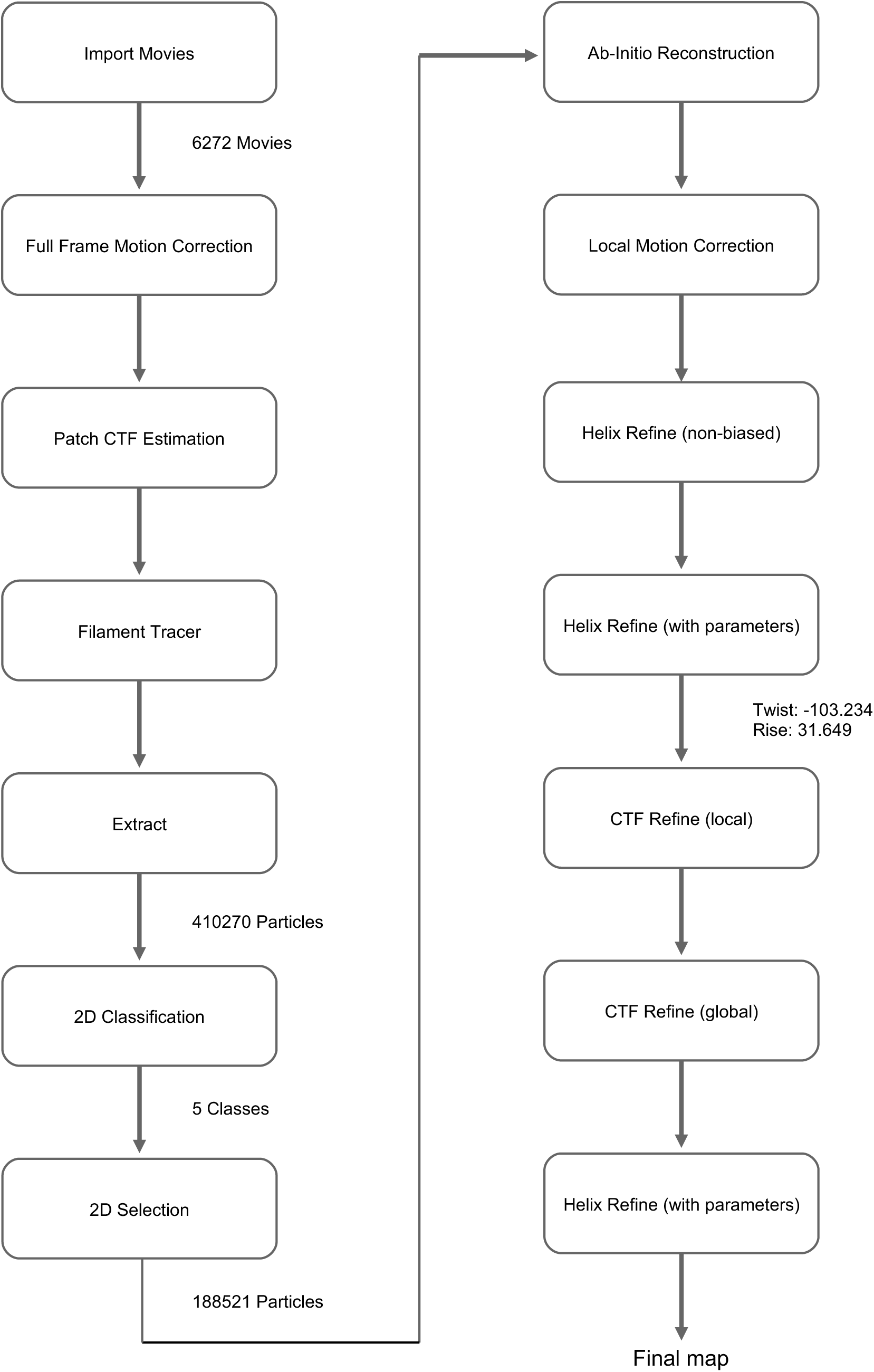
Image processing pipeline. Flowchart showing the helical reconstruction workflow in cryoSPARC

**Supplementary Figure 4.**
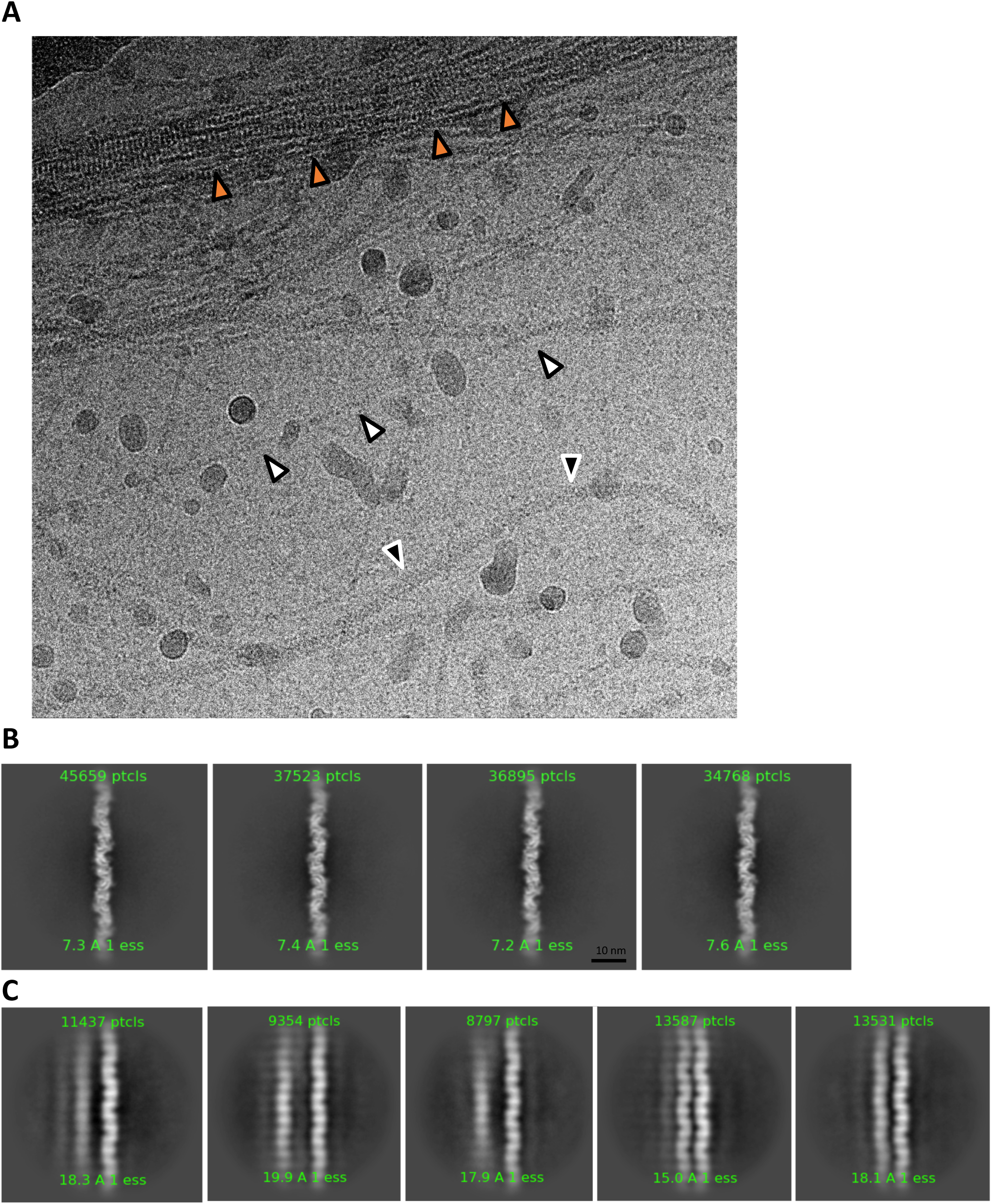
CryoEM and 2D classification. **A,** cryoEM micrograph showing a mixture of thread and AAP filaments from *S. acidocaldarius*. White arrowheads, Threads; orange arrowheads, cables of Threads; black arrowheads, AAP or archaella. **B,** 2D classification of thread filaments carried out in cryoSPARC. **C,** 2D classification showing the tendency of threads to line up in parallel cables.

**Supplementary Figure 5.**
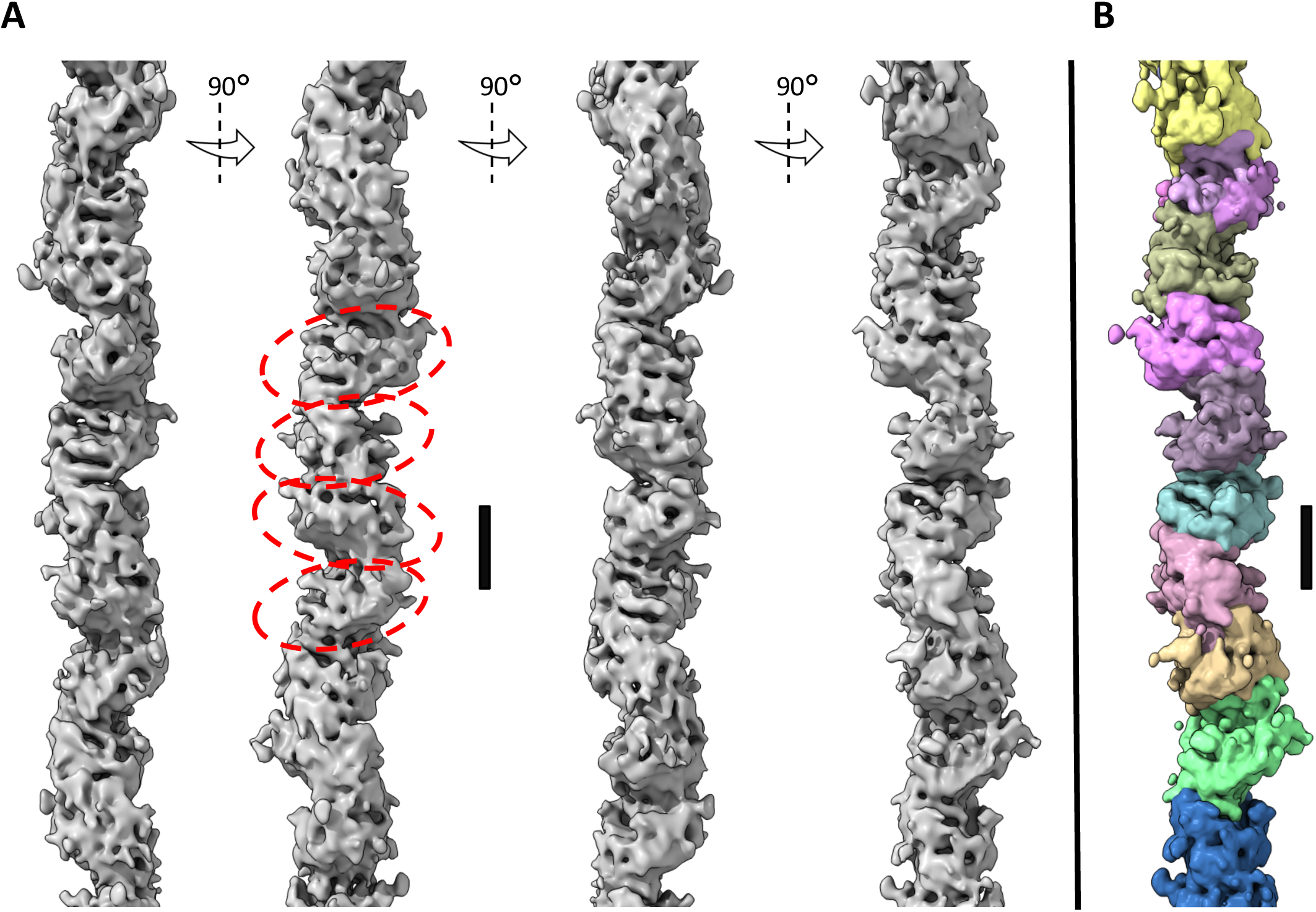
3D reconstruction without helical parameters. **A**, Unbiased helix refinement generated in cryoSPARC without the input of helical parameters. Careful inspection of the density indicates that the subunit rise is approximately 30 Å (black scale bar). Individual subunits are indicated with dashed ovals. **B**, segmented map showing the filament’s subunits in different colours.

**Supplementary Figure 6.**
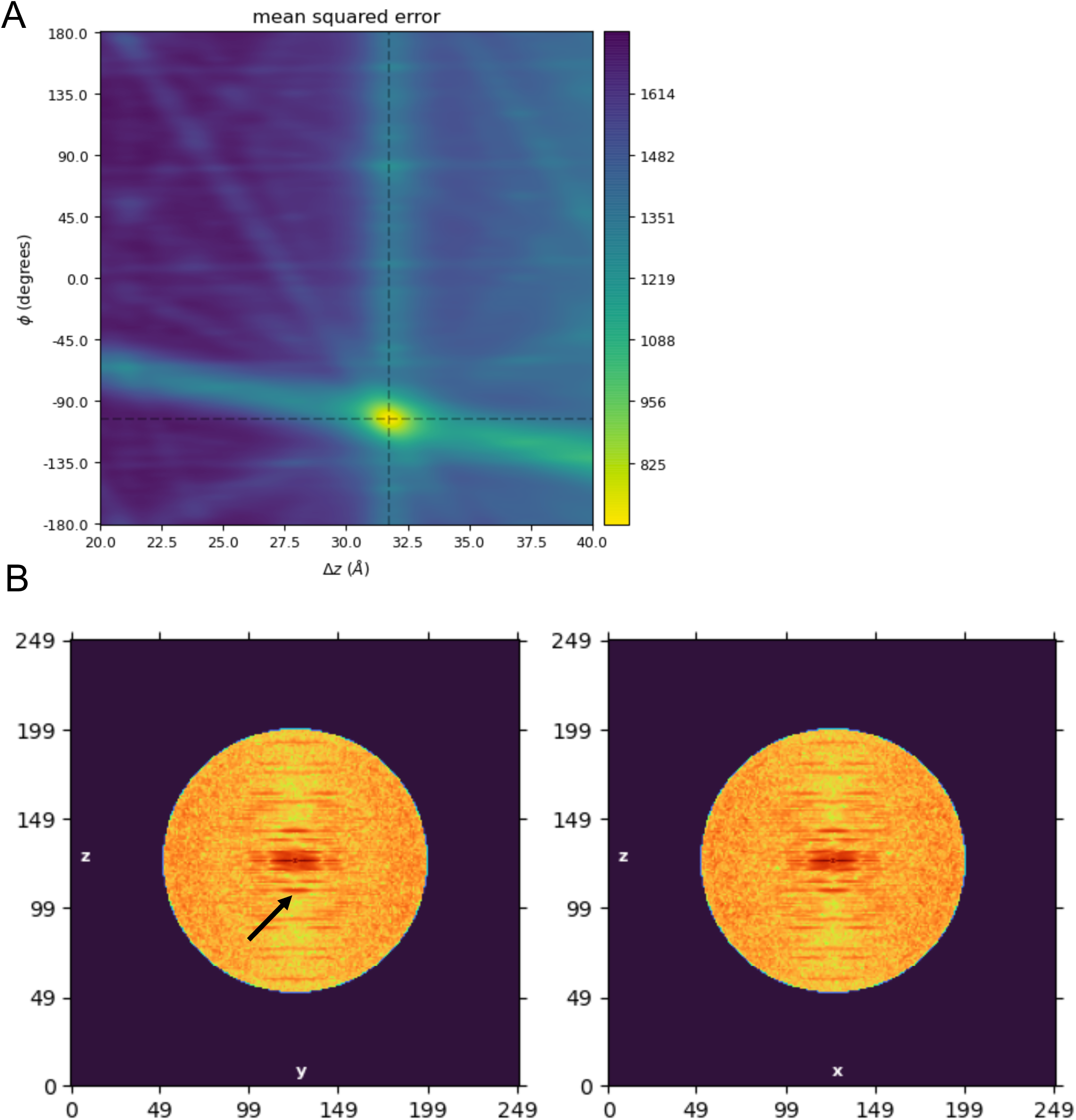
Helical parameter search. **A,** cryoSPARC symmetry search showing a clear peak at 31.6 Å rise and -103° twist. **B**, The helical rise is confirmed by a meridomal reflection in the power spectrum of the Thread filament, which corresponds to a value of 31.4 Å.

**Supplementary Figure 7.**
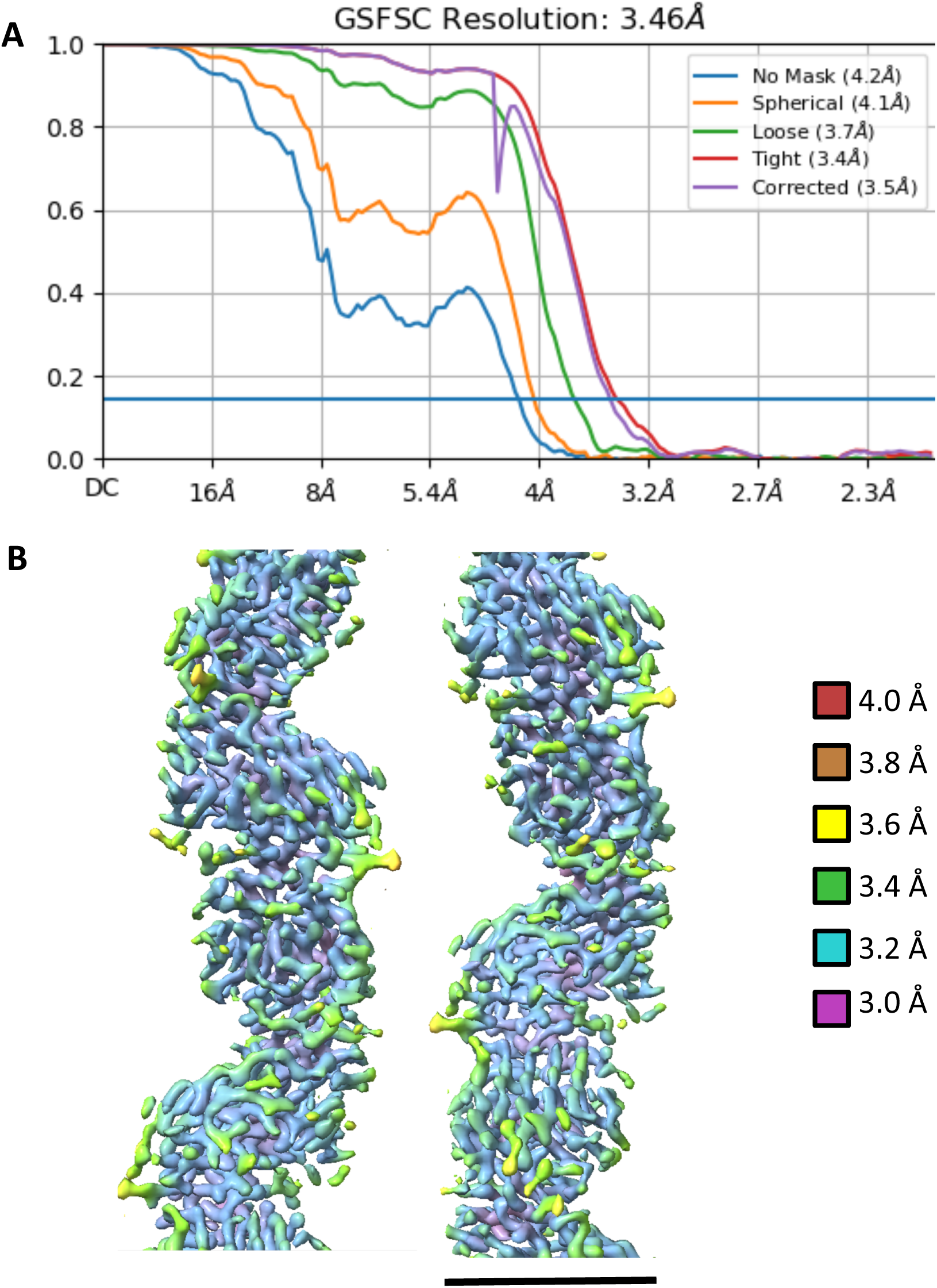
Global and local resolution estimation. **A,** Global resolution of the helical reconstruction of the thread filament. **B,** Local resolution map shows a resolution range of ∼3 – 4 Å. The core of the filament is resolved best, while peripheral regions are slightly less well defined. This is particularly the case for the flexible glycan moieties. Scale 40 Å

**Supplementary Figure 8.**
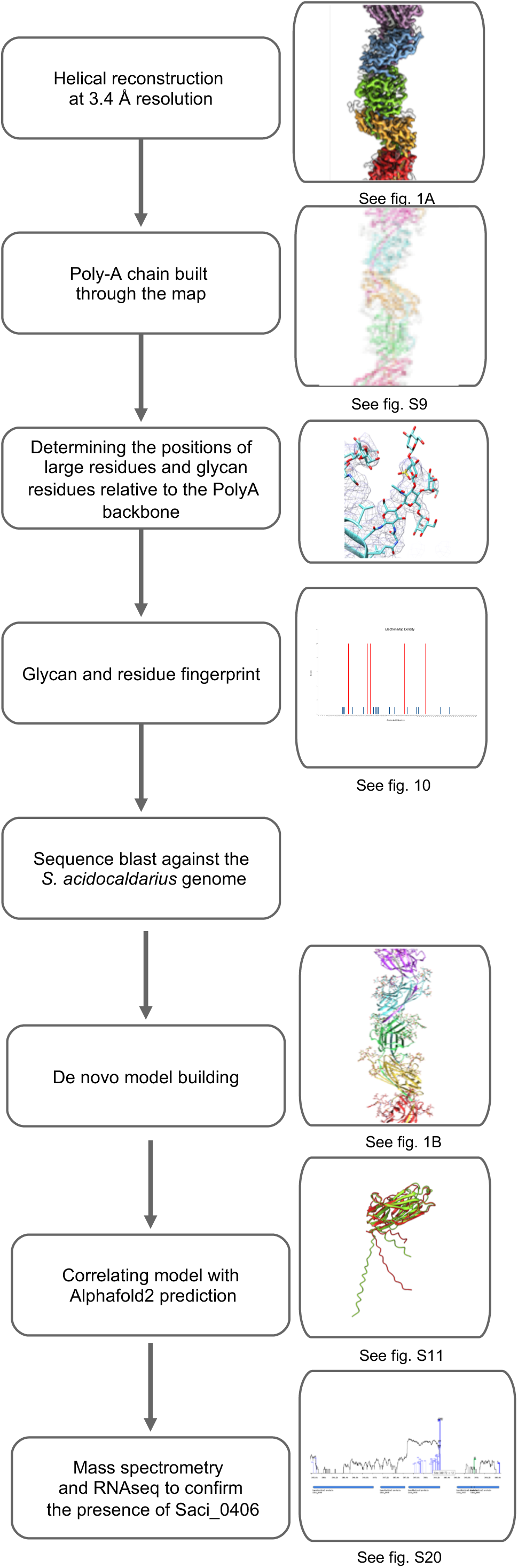
Determining the identity of the thread subunit protein from the cryoEM map. Flow chart illustrating how the identity of the thread subunit Saci_0406 was revealed based on the cryoEM map

**Supplementary Figure 9.**
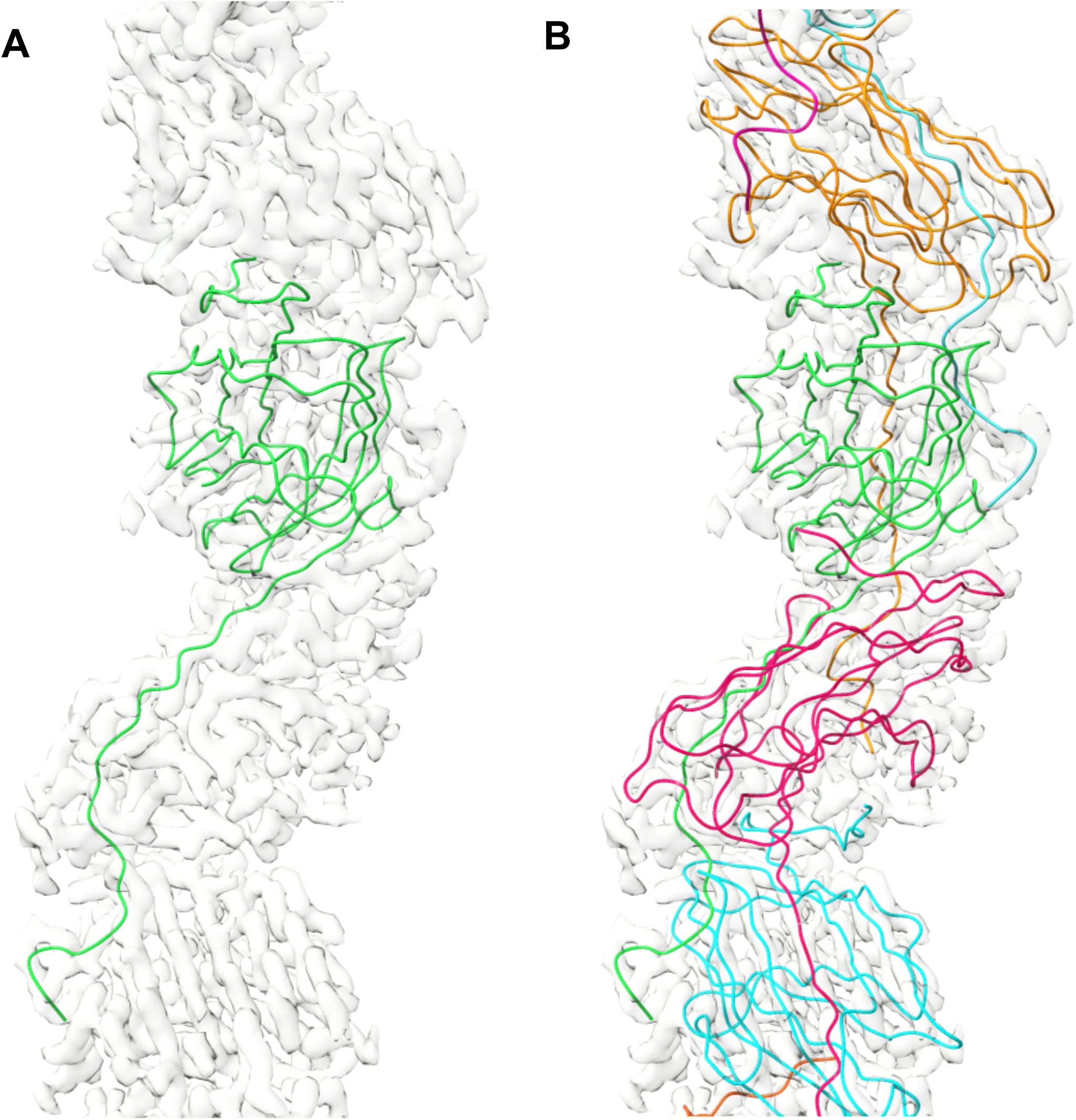
Poly alanine model. **A**, A poly-alanine chain built into the density of one monomer. **B,** Four poly alanine chains corresponding to four subunits built into the cryoEM map.

**Supplementary Figure 10.**
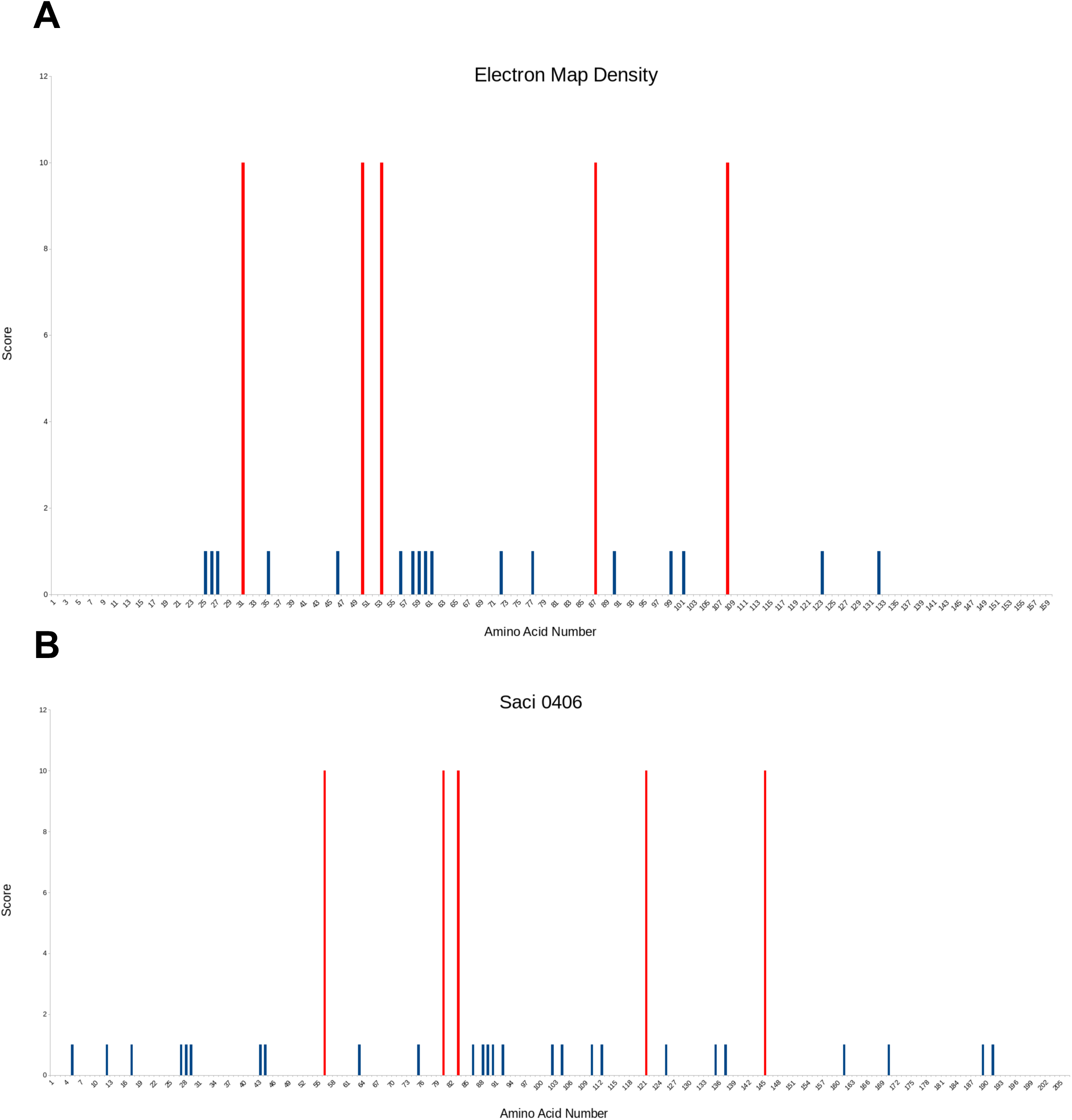
Glycan and large residue fingerprint. **A,** Bar chart illustrating the positions of large side chains and putative glycan densities relative to a poly-alanine chain treaded though the cryoEM map. For illustration purposes, glycans were given a value of 10 (red bars) and visibly aromatic amino acid residues a value of 1 (blue bars). **B,** bar chart illustrating the position of conserved glycosylation sequons (**N**XS/T) (red) and aromatic residues (blue) within the sequence of Saci_0406.

**Supplementary Figure 11.**
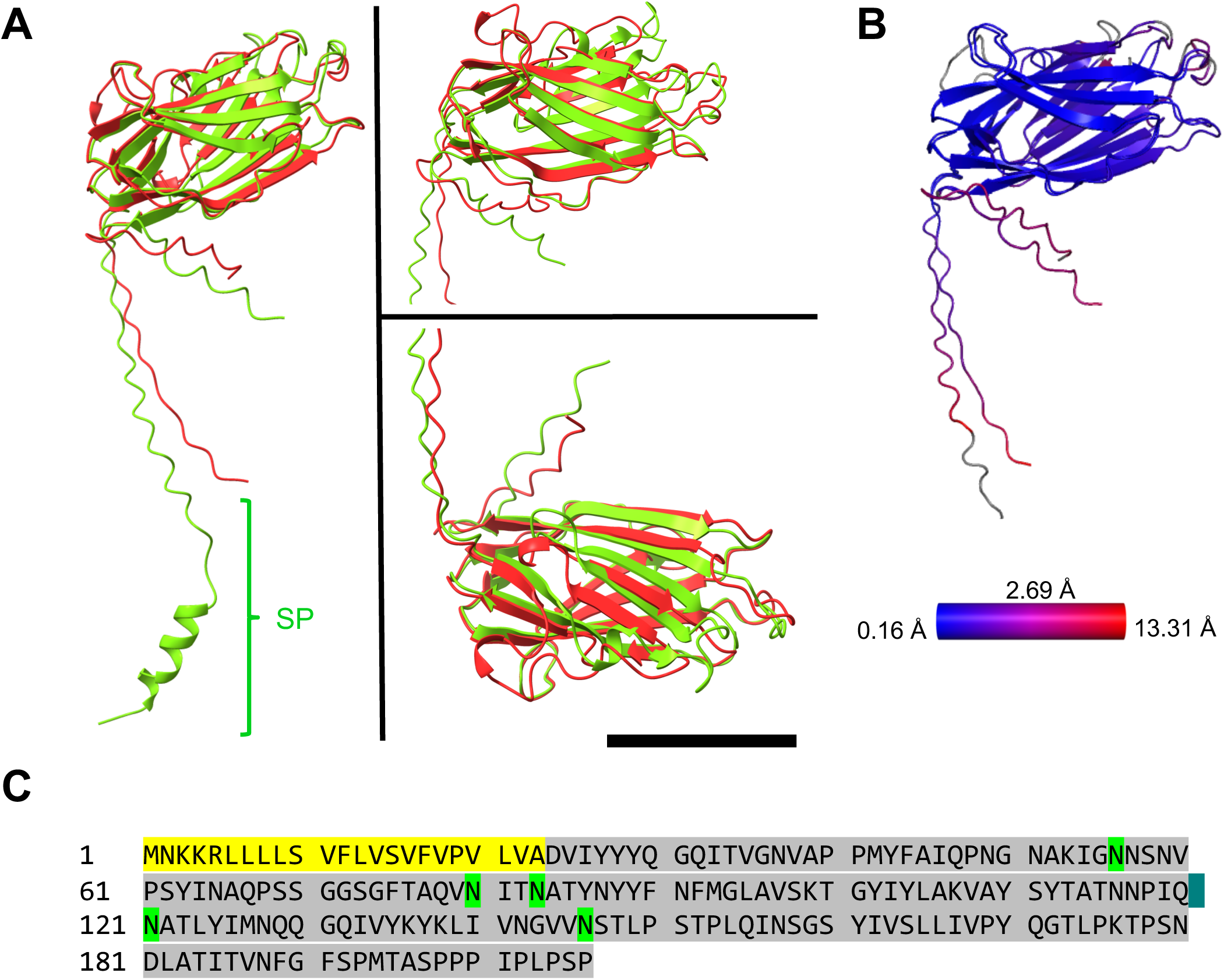
*Ab initio* structure vs Alphafold2 prediction and sequence of Saci_0406. **A,** *ab initio* atomic model of Saci_0408 (red) superimposed with the Alphafold2 prediction of the protein (green). Note that the Alphafold2 prediction additionally contains the N-terminal signal peptide (SP). **B,** RMSD values between the two structures in A (blue, low; red high RMSD). Average value 2.69 Å. **C,** Protein sequence of Saci_0406 with the cleaved signal sequence highlighted in yellow and glycosylation sites in green. Each of the glycosylated asparagine residues resides in a NXS/T sequon. Scale bar 20 Å

**Supplementary Figure 12.**
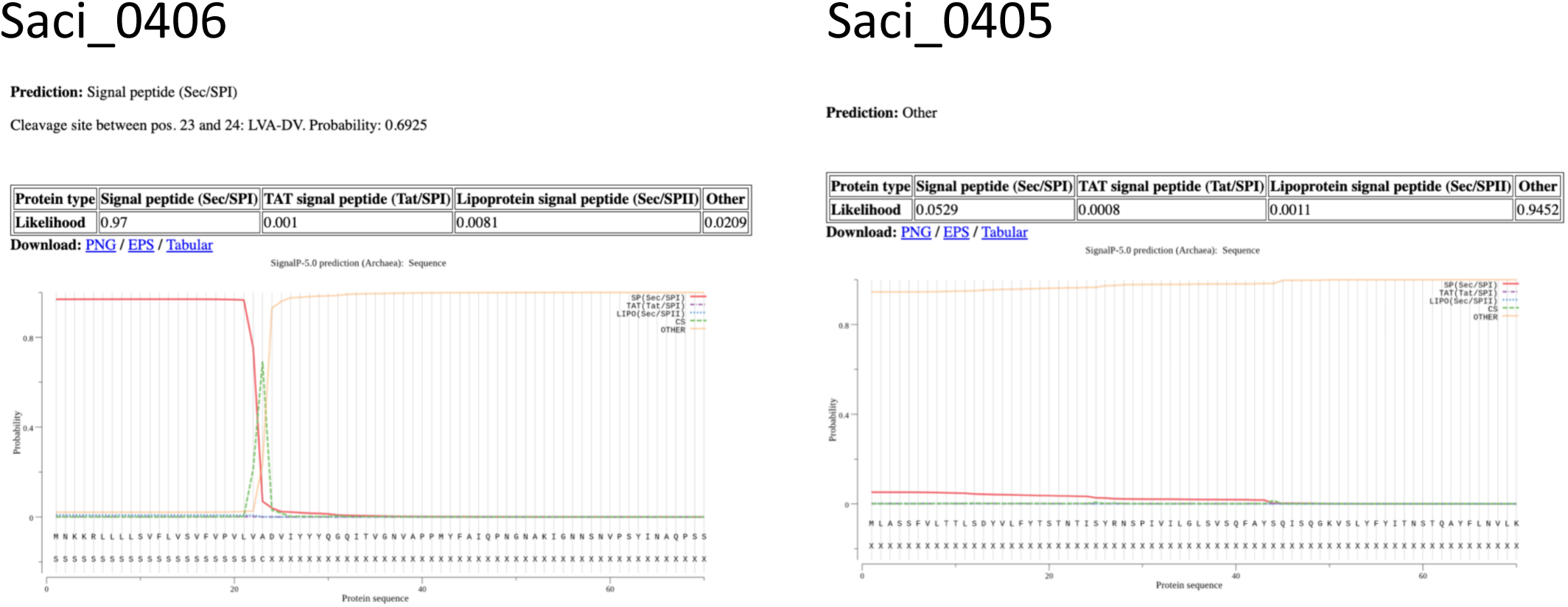
Signal sequence prediction using the SignalP 5.0 server. **Left**, prediction for Saci_0406 indicates a SEC signal sequence cleaved by signal peptidase 1 (SP1). **Right**, prediction for Saci_0405 shows no SEC signal sequence.

**Supplementary Figure 13.**
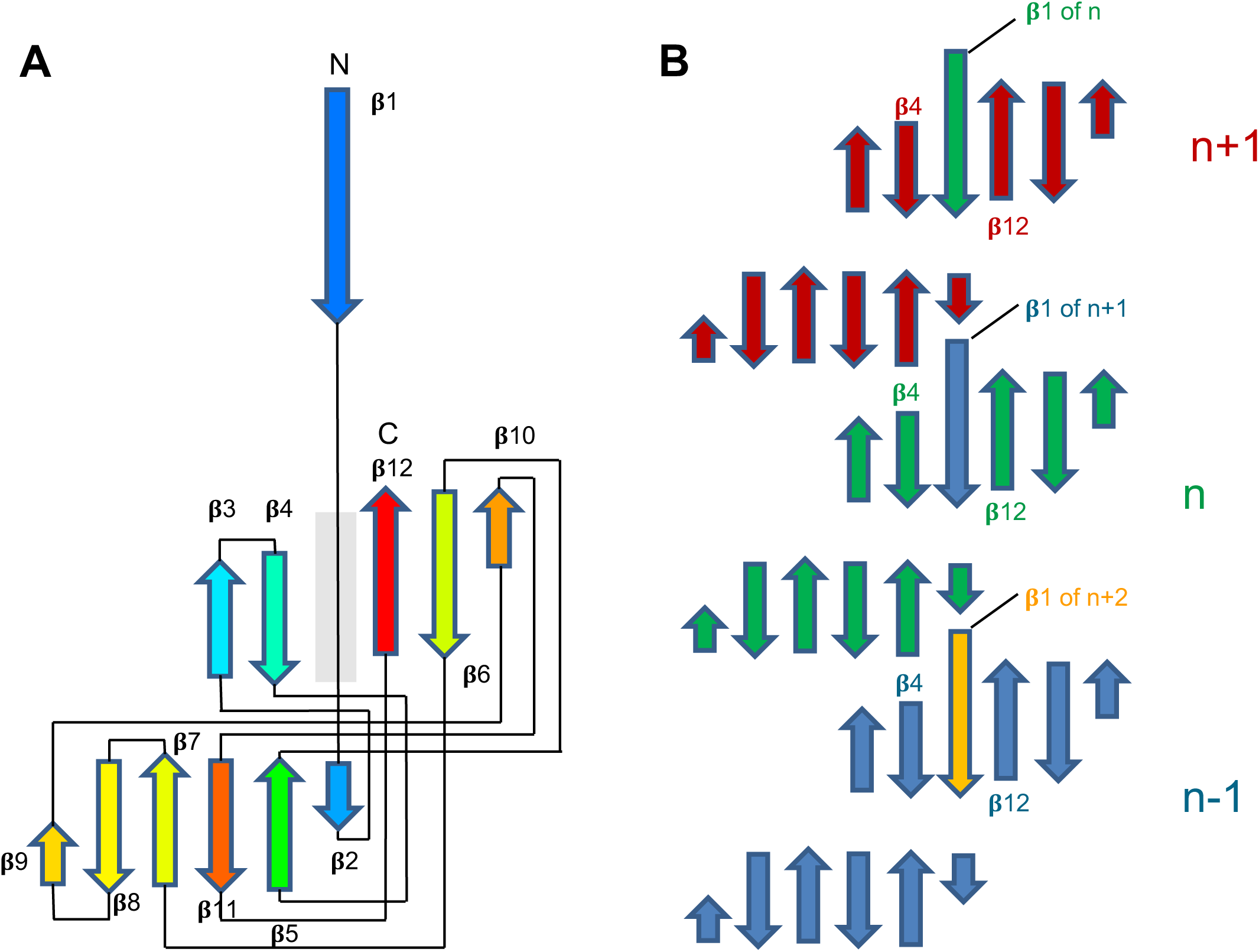
Topology diagram of the thread. **A,** Topology of Saci_0406 in rainbow colouring (blue, N-terminus; red, C-terminus). The subunit consists of 12 𝛃 strands (numbered). A gap between 𝛃4 and 𝛃12 serves as 𝛃-strand acceptor site (transparent grey box). 𝛃1 acts as 𝛃-strand donor for an adjacent monomer. **B**, topology of three Saci_0406 subunits within the thread. The central monomer is named n. n-1 is the monomer preceding n and n+1 is the monomer following n. The monomer n inserts its 𝛃-strand into the acceptor site between 𝛃4 and 𝛃12 in the monomer n+1. The monomer n+1 inserts its 𝛃-strand into the acceptor site between 𝛃4 and 𝛃12 in the monomer n+2 and so forth.

**Supplementary Figure 14.**
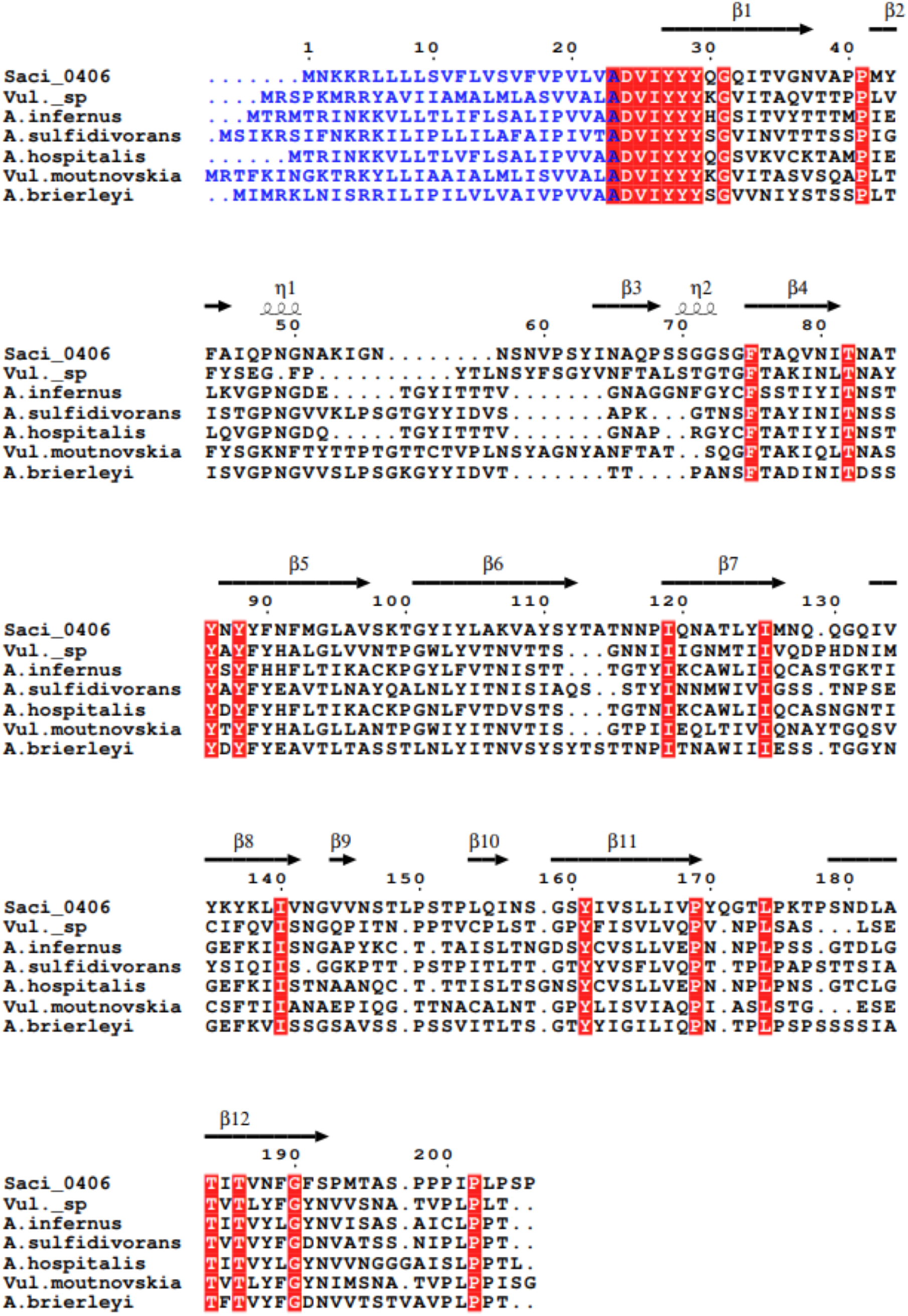
Multisequence alignment between Saci_0406 homologs in related archaeal species.

**Supplementary Figure 15.**
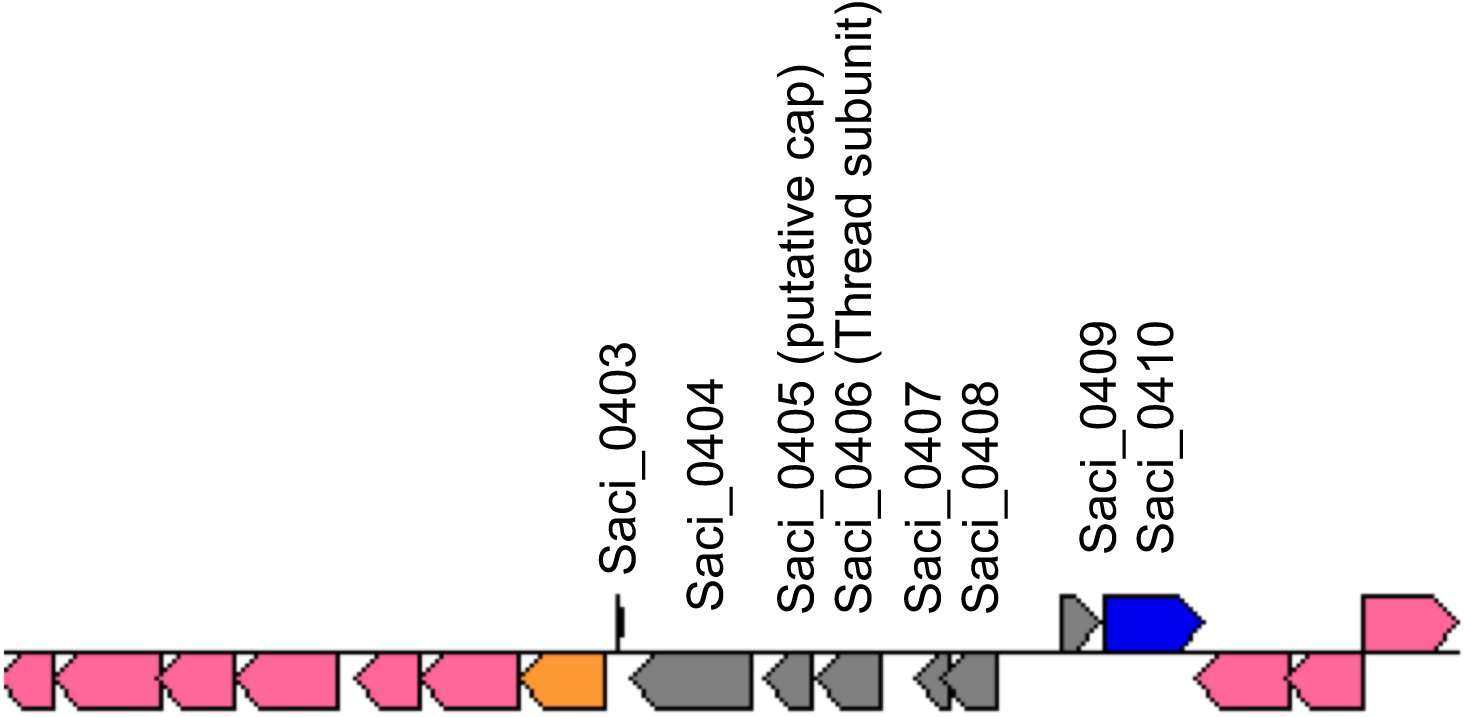
Gene cluster surrounding Saci_0406. Saci_0406 encodes for the thread monomer. Saci_0405 encodes for a hypothetical cap protein. Saci_0407 and Saci_0408 may form parts of the assembly machinery.

**Supplementary Figure 16.**
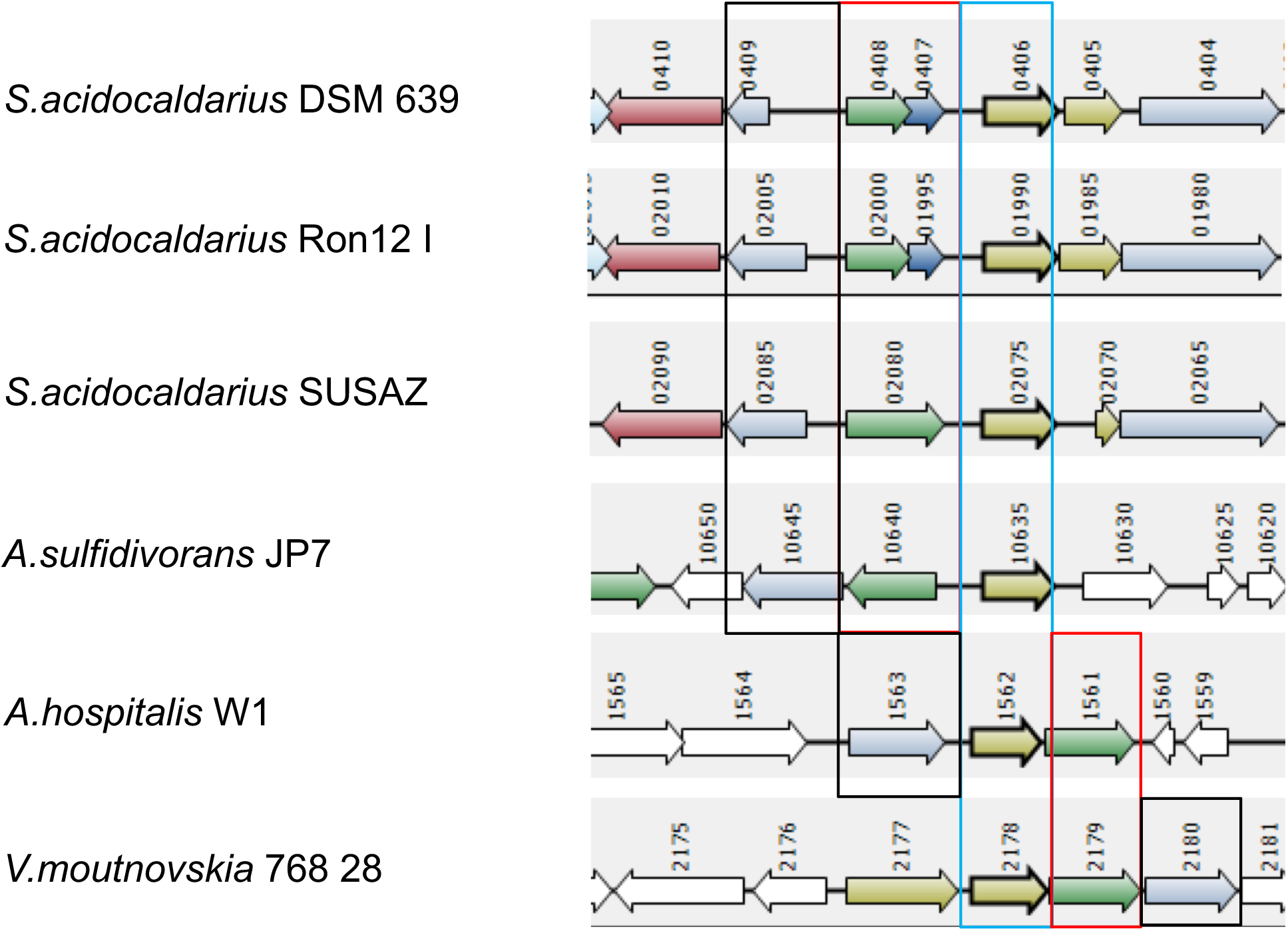
The thread gene cluster is found in various *S.aci* strains. **A,** diagram visualising the similarities between genes in different strains of *S. acidocaldarius* and other related species. Red box – homologues of *Saci_0407* and *Saci_0408*. In the *acidocaldarius* strains DSM 639 and Ron12, *Saci_0407* and *Saci_0408* are two separate genes. In the *S.aci* SUSAZ strain, the *Saci_0407* and *Saci_0408* homologs are fused into one gene, *SUSAZ_02080*. Blue box – thread subunit gene (*Saci_0406*) homologues. Black box – Saci_0409 homologues in the various strains and species. The length of Saci_0409 seems to be variable between different organisms, but the gene is always present.

**Supplementary Figure 17.**
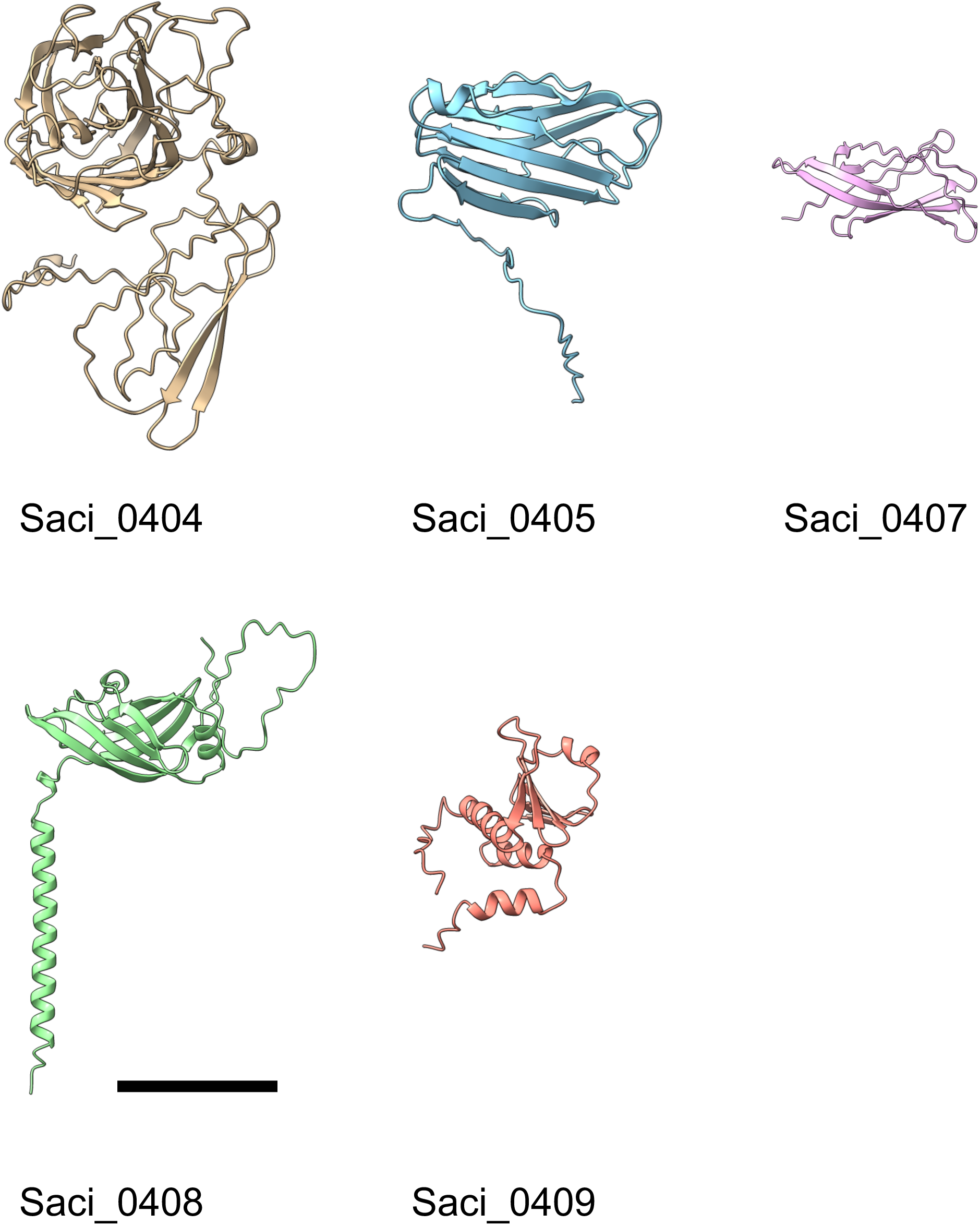
Gallery of structural predictions for the proteins encoded in the proposed operon surrounding *saci_0406*. Structures were predicted using Alphafold2. Scale bar 20 Å

**Supplementary Figure 18.**
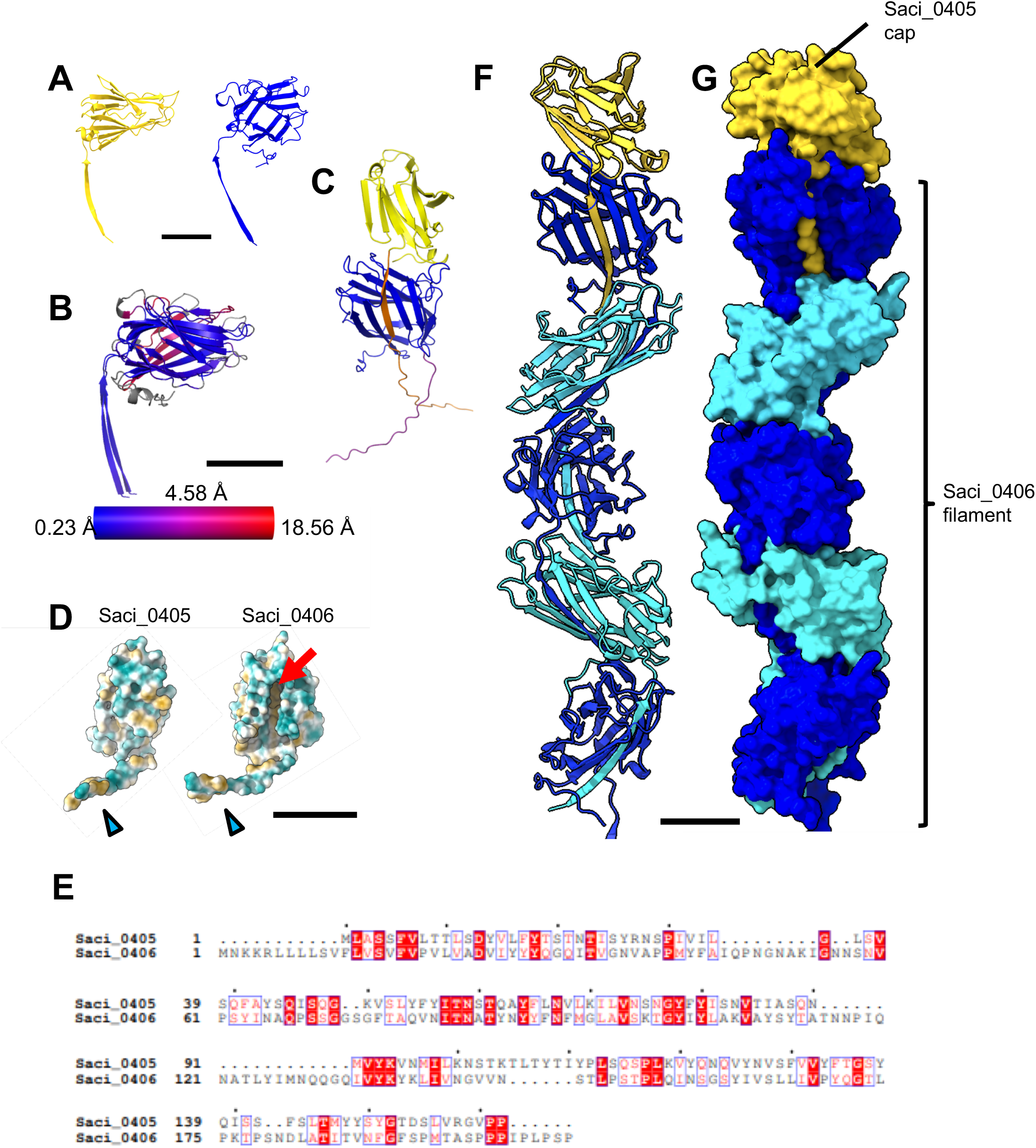
The putative cap protein Saci_0405. **A**, Alphafold2 prediction of Saci_0405 (yellow) compared with the experimentally determined structure of the thread subunit Saci_0406 (blue). **B**, alignment of the models for Saci_0405 and Saci_0406 with RMSD value coloured from blue (lowest) to red (highest), highlighting the structural similarity between of both proteins. The average RMSD is 4.58 Å. **C,** Alphafold2 model of the Saci_0406 (blue) and Saci_0405 (yellow) complex predicts donor strand complementation (orange). **D**, Saci_0405 and Saci_0406 shown in solid representation and coloured by hydrophobicity (orange hydrophobic, blue, hydrophilic). Both proteins have conserved tail domains that can act as beta-strand donors (blue arrowheads). However, Saci_0405 lacks the acceptor groove for a tail domain of a subsequent monomer, which is present in Saci_0406 (red arrow). This suggests that Saci_0405 could act as a terminal cap protein. **E,** sequence alignment between Saci_0405 and Saci_0406. **F, G,** proposed filament structure of Saci_0406 monomers (blue) with Saci_0405 modelled as the terminal cap in ribbon (E) and solid representation (F). Glycans have been omitted for simplicity. Scale bar 20 Å

**Supplementary Figure 19.**
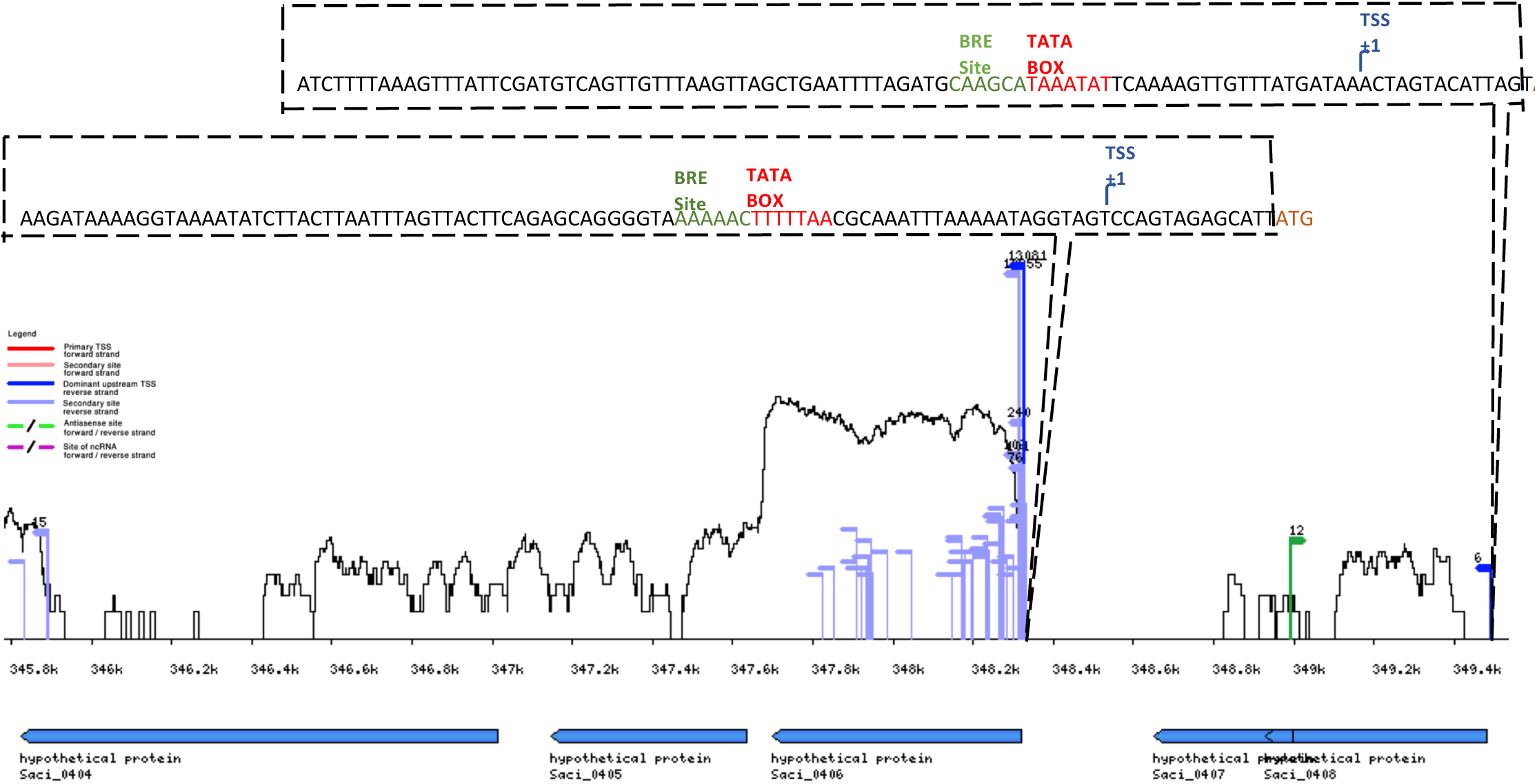
Transcriptomics profile of the proposed thread operon.

**Supplementary Figure 20.**
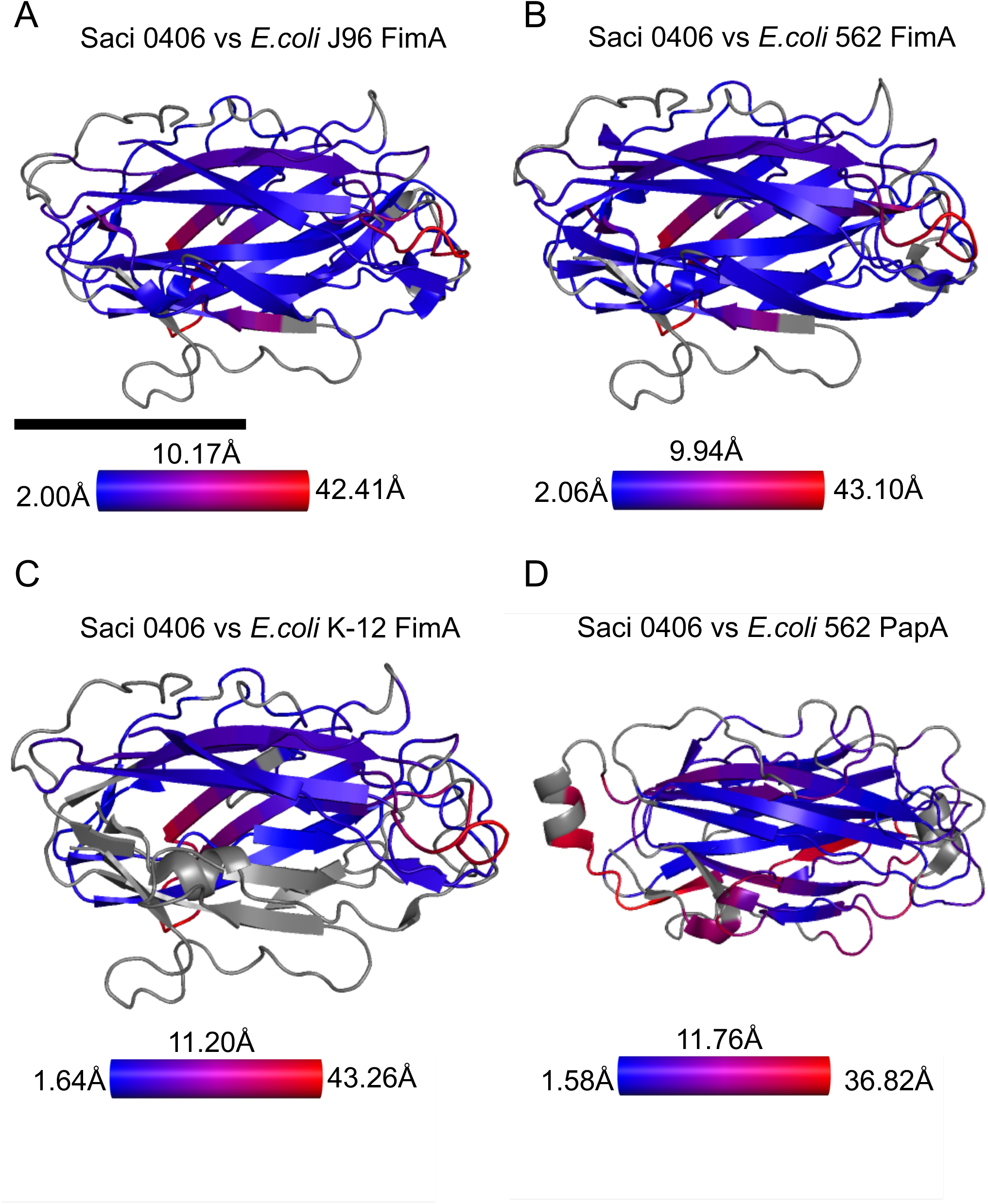
Structural similarity between Saci_0406 and known *E.coli* chaperone-usher pilins. **A - D** RMSD values calculated for pairs of Saci_0406 and *E.coli* chaperone-usher pili, where blue indicates low values, red high, and grey areas ignored. **A,** Average RMSD 10.17Å, minimum value 2.00Å, maximum value 42.41Å. **B,** Average RMSD 9.94Å, minimum value 2.06Å, maximum value 43.10Å. **C,** Average RMSD 11.20Å, minimum value 1.64Å, maximum value 43.26Å. **D,** Average RMSD 11.76Å, minimum value 1.58Å, maximum value 36.82Å. Scale bar 20Å

**Supplementary Figure 21.**
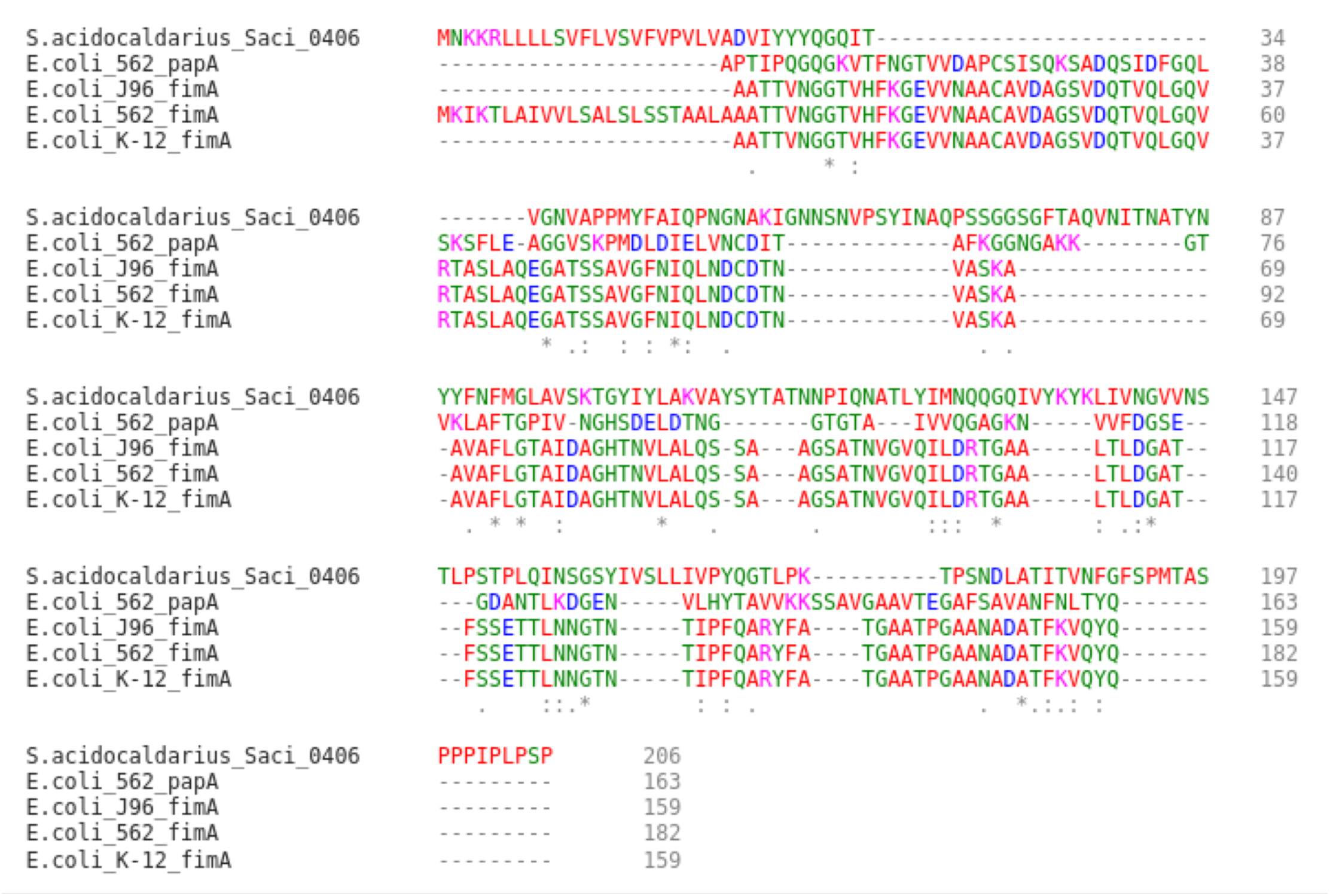
Multisequence alignment of Saci_0406 and known *E.coli* type-I chaperone-usher pili.

**Supplementary Figure 22.**
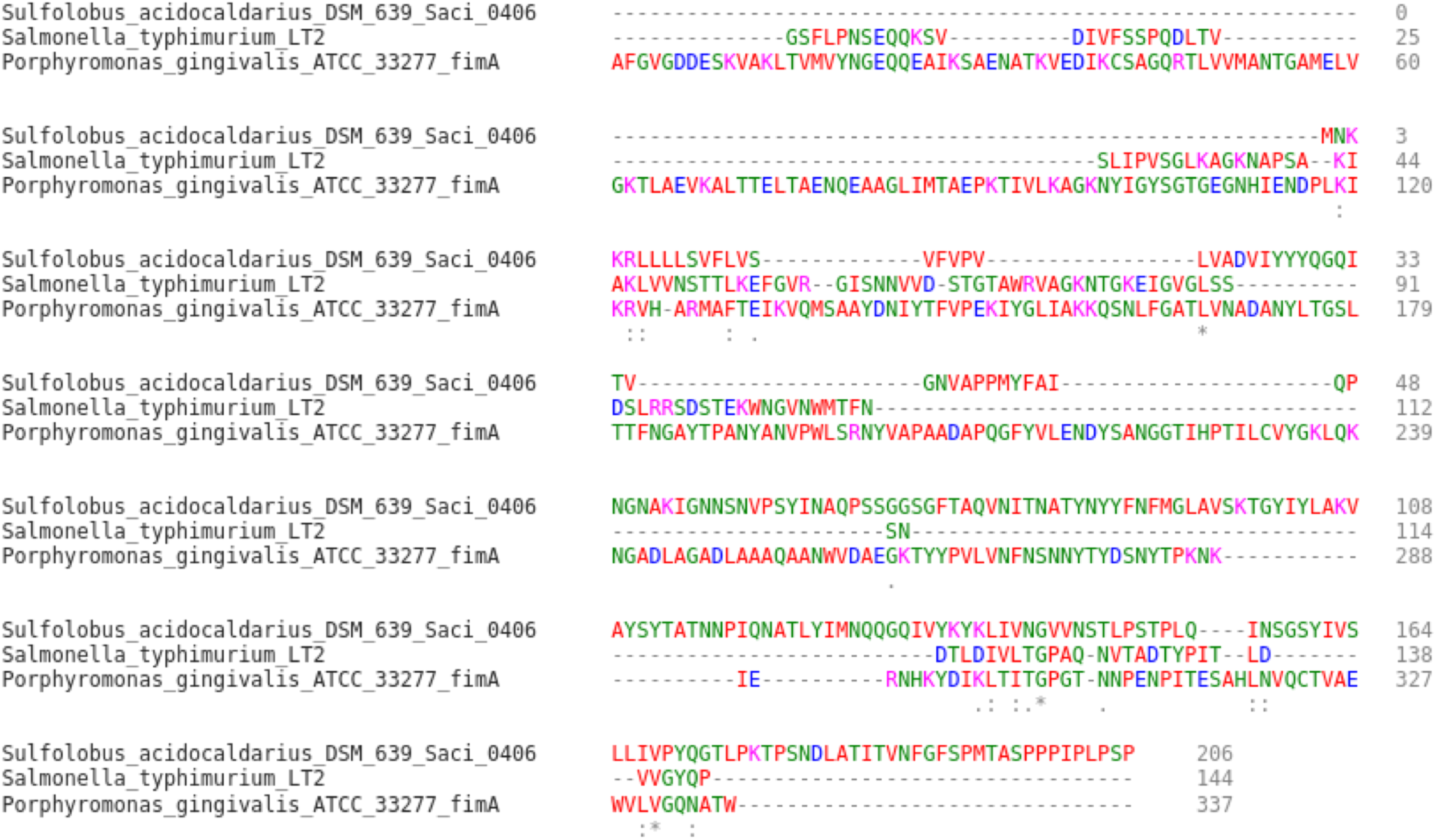
Multisequence alignment of Saci_0406 and subunits of type-V pili (SafA and FimA).

## Notes

### Competing Interest Statement

The authors have declared no competing interest.

## References

1. Costa, T. R. D. et al. Secretion systems in Gram-negative bacteria: structural and mechanistic insights. Nat. Rev. Microbiol. 13, 343–359 (2015).

2. Hospenthal, M. K., Costa, T. R. D. & Waksman, G. A comprehensive guide to pilus biogenesis in Gram-negative bacteria. Nat. Rev. Microbiol. 15, 365–379 (2017).

3. Meyer, B. H. & Albers, S.-V. Hot and sweet: protein glycosylation in Crenarchaeota. Biochem. Soc. Trans. 41, 384–392 (2013).

4. Makarova, K. S., Koonin, E. V. & Albers, S.-V. V. Diversity and Evolution of Type IV pili Systems in Archaea. Front. Microbiol. 7, 667 (2016).

5. Pohlschroder, M. & Esquivel, R. N. Archaeal type IV pili and their involvement in biofilm formation. Front. Microbiol. 6, 190 (2015).

6. Denise, R., Abby, S. S. & Rocha, E. P. C. C. Diversification of the type IV filament superfamily into machines for adhesion, protein secretion, DNA uptake, and motility. PLoS Biol. 17, e3000390 (2019).

7. Brock, T. D., Brock, K. M., Belly, R. T. & Weiss, R. L. Sulfolobus: A new genus of sulfur- oxidizing bacteria living at low pH and high temperature. Arch. für Mikrobiol. 1972 841 84, 54–68 (1972).

8. Jarrell, K. F., Albers, S.-V. & Machado, J. N. de S. A comprehensive history of motility and Archaellation in Archaea. FEMS Microbes 2, 2 (2021).

9. Beeby, M., Ferreira, J. L., Tripp, P., Albers, S.-V. & Mitchell, D. R. Propulsive nanomachines: the convergent evolution of archaella, flagella and cilia. FEMS Microbiol. Rev. 44, 253–304 (2020).

10. Bardy, S. L. & Jarrell, K. F. FlaK of the archaeon Methanococcus maripaludis possesses preflagellin peptidase activity. FEMS Microbiol. Lett. 208, 53–9 (2002).

11. Albers, S.-V., Szabó, Z. & Driessen, A. J. M. Archaeal Homolog of Bacterial Type IV Prepilin Signal Peptidases with Broad Substrate Specificity. J. Bacteriol. 185, 3918–3925 (2003).

12. Poweleit, N. et al. CryoEM structure of the Methanospirillum hungatei archaellum reveals structural features distinct from the bacterial flagellum and type IV pilus. Nat. Microbiol. 2, 16222 (2017).

13. Jarrell, et al. N-Linked Glycosylation in Archaea: a Structural, Functional, and Genetic Analysis. Microbiol. Mol. Biol. Rev. 78, 304–341 (2014).

14. Daum, B. et al. Structure and in situ organisation of the Pyrococcus furiosus archaellum machinery. Elife 6, (2017).

15. Meshcheryakov, V. A. et al. High-resolution archaellum structure reveals a conserved metal-binding site. EMBO Rep. 20, (2019).

16. Gambelli, L. et al. An archaellum filament composed of two alternating subunits. Nat. Commun. 2022 131 13, 1–11 (2022).

17. Albers, S.-V. & Meyer, B. H. The archaeal cell envelope. Nat. Rev. Microbiol. 9, 414–26 (2011).

18. Wang, F. et al. The structures of two archaeal type IV pili illuminate evolutionary relationships. Nat. Commun. 2020 111 11, 1–10 (2020).

19. Wang, F. et al. An extensively glycosylated archaeal pilus survives extreme conditions. Nat. Microbiol. 4, 1401–1410 (2019).

20. Hartman, R. et al. The Molecular Mechanism of Cellular Attachment for an Archaeal Virus. Structure 27, 1634–1646.e3 (2019).

21. Rowland, E. F., Bautista, M. A., Zhang, C. & Whitaker, R. J. Surface resistance to SSVs and SIRVs in pilin deletions of *Sulfolobus islandicus*. Mol. Microbiol. 113, 718–727 (2020).

22. van Wolferen, M. et al. Species-Specific Recognition of Sulfolobales Mediated by UV- Inducible Pili and S-Layer Glycosylation Patterns. MBio 11, (2020).

23. van Wolferen, M., Ajon, M., Driessen, A. J. M. & Albers, S.-V. Molecular analysis of the UV-inducible pili operon from Sulfolobus acidocaldarius. Microbiologyopen 2, 928–937 (2013).

24. van Wolferen, M., Ajon, M., Driessen, A. J. M. & Albers, S.-V. Molecular analysis of the UV-inducible pili operon from *Sulfolobus acidocaldarius*. Microbiologyopen 2, 928–937 (2013).

25. Van Wolferen, M., Wagner, A., Van Der Does, C. & Albers, S. V. The archaeal Ced system imports DNA. Proc. Natl. Acad. Sci. U. S. A. 113, 2496–2501 (2016).

26. van Wolferen, M., Ajon, M., Driessen, A. J. M. & Albers, S.-V. How hyperthermophiles adapt to change their lives: DNA exchange in extreme conditions. Extremophiles 17, 545–563 (2013).

27. Henche, A.-L. et al. Structure and function of the adhesive type IV pilus of Sulfolobus acidocaldarius. Environ. Microbiol. 14, 3188–3202 (2012).

28. Henche, A.-L., Koerdt, A., Ghosh, A. & Albers, S.-V. V. Influence of cell surface structures on crenarchaeal biofilm formation using a thermostable green fluorescent protein. Environ. Microbiol. 14, 779–93 (2012).

29. Punjani, A., Rubinstein, J. L., Fleet, D. J. & Brubaker, M. A. CryoSPARC: Algorithms for rapid unsupervised cryo-EM structure determination. Nat. Methods 14, 290–296 (2017).

30. Desfosses, A., Ciuffa, R., Gutsche, I. & Sachse, C. SPRING - an image processing package for single-particle based helical reconstruction from electron cryomicrographs. J. Struct. Biol. 185, 15–26 (2014).

31. Egelman, E. H. et al. Structural plasticity of helical nanotubes based on coiled-coil assemblies. Structure 23, 280–289 (2015).

32. Kolappan, S., Tracy, E. N., Bakaletz, L. O., Munson, R. S. & Craig, L. Expression, purification, crystallization and preliminary crystallographic analysis of PilA from the nontypeable Haemophilus influenzae type IV pilus. Acta Crystallogr. Sect. F. Struct. Biol. Cryst. Commun. 68, 284–287 (2012).

33. Almagro Armenteros, J. J., et al. SignalP 5.0 improves signal peptide predictions using deep neural networks. Nat. Biotechnol. 37, 420–423 (2019).

34. Shibata, S. et al. Structure of polymerized type V pilin reveals assembly mechanism involving protease-mediated strand exchange. Nat. Microbiol. 5, 830–837 (2020).

35. Salih, O., Remaut, H., Waksman, G. & Orlova, E. V. Structural Analysis of the Saf Pilus by Electron Microscopy and Image Processing. J. Mol. Biol. 379, 174–187 (2008).

36. Meyer, B. H. & Albers, S.-V. Hot and sweet: protein glycosylation in Crenarchaeota. Biochem. Soc. Trans. 41, (2013).

37. Peyfoon, E. et al. The S-layer glycoprotein of the crenarchaeote Sulfolobus acidocaldarius is glycosylated at multiple sites with chitobiose-linked N-glycans. Archaea 2010, (2010).

38. Huerta-Cepas, J. et al. eggNOG 5.0: a hierarchical, functionally and phylogenetically annotated orthology resource based on 5090 organisms and 2502 viruses. Nucleic Acids Res. 47, D309–D314 (2019).

39. Wurtzel, O. et al. A single-base resolution map of an archaeal transcriptome. Genome Res. 20, 133–141 (2010).

40. Zeng, L., Zhang, L., Wang, P. & Meng, G. Structural basis of host recognition and biofilm formation by Salmonella Saf pili. Elife 6, (2017).

41. Hospenthal, M. K. et al. The cryoelectron microscopy structure of the Type 1 Chaperone-Usher pilus rod. Structure 25, 1829–1838.e4 (2017).

42. Hospenthal, M. K. et al. Structure of a Chaperone-Usher Pilus reveals the molecular basis of rod uncoiling. Cell 164, 269–278 (2016).

43. Du, M. et al. Processive dynamics of the usher assembly platform during uropathogenic Escherichia coli P pilus biogenesis. Nat. Commun. 12, (2021).

44. Omattage, N. S. et al. Structural basis for usher activation and intramolecular subunit transfer in P pilus biogenesis in Escherichia coli. Nat. Microbiol. 3, 1362–1368 (2018).

45. Alonso-Caballero, A. et al. Mechanical architecture and folding of E. coli type 1 pilus domains. Nat. Commun. 9, (2018).

46. Wang, F., Cvirkaite-Krupovic, V., Krupovic, M. & Egelman, E. H. Archaeal bundling pili of Pyrobaculum calidifontis reveal similarities between archaeal and bacterial biofilms. bioRxiv 2022.04.22.489182 (2022) doi:10.1101/2022.04.22.489182.

47. Hagan, R. M., et al. NMR spectroscopic and theoretical analysis of a spontaneously formed lys-asp isopeptide bond. Angew. Chemie - Int. Ed. 49, 8421–8425 (2010).

48. Hendrickx, A. P. A. et al. Isopeptide bonds of the major pilin protein BcpA influence pilus structure and bundle formation on the surface of Bacillus cereus. Mol. Microbiol. 85, 152–163 (2012).

49. Kang, H. J. & Baker, E. N. Intramolecular isopeptide bonds give thermodynamic and proteolytic stability to the major Pilin protein of Streptococcus pyogenes. J. Biol. Chem. 284, 20729–20737 (2009).

50. Persson, K., Esberg, A., Claesson, R. & Strömberg, N. The Pilin Protein FimP from Actinomyces oris: Crystal Structure and Sequence Analyses. PLoS One 7, (2012).

51. Hae, J. K., Coulibaly, F., Clow, F., Proft, T. & Baker, E. N. Stabilizing isopeptide bonds revealed in gram-positive bacterial pilus structure. Science (80-.). 318, 1625–1628 (2007).

52. Abdul Halim, M. F., et al. Haloferax volcanii archaeosortase is required for motility, mating, and C-terminal processing of the S-layer glycoprotein. Mol. Microbiol. 88, 1164–1175 (2013).

53. Siegmund, V. et al. Spontaneous Isopeptide bond formation as a powerful tool for engineering Site-Specific antibody-drug conjugates. Sci. Reports 2016 61 6, 1–9 (2016).

54. Meyer, B. H., Birich, A. & Albers, S. V. N-Glycosylation of the archaellum filament is not important for archaella assembly and motility, although N-Glycosylation is essential for motility in Sulfolobus acidocaldarius. Biochimie 118, 294–301 (2015).

55. Nielsen, H., Brunak, S. & Von Heijne, G. Machine learning approaches for the prediction of signal peptides and other protein sorting signals. Protein Eng. Des. Sel. 12, 3–9 (1999).

56. Meyer, B. H. & Albers, S.-V. AglB, catalyzing the oligosaccharyl transferase step of the archaeal N-glycosylation process, is essential in the thermoacidophilic crenarchaeon Sulfolobus acidocaldarius. Microbiologyopen 3, 531–543 (2014).

57. Vetsch, M. et al. Pilus chaperones represent a new type of protein-folding catalyst. Nat. 2004 4317006 431, 329–333 (2004).

58. Sauer, F. G. et al. Structural Basis of Chaperone Function and Pilus Biogenesis. Science (80-.). 285, 1058–1061 (1999).

59. Sauer, F. G., Pinkner, J. S., Waksman, G. & Hultgren, S. J. Chaperone Priming of Pilus Subunits Facilitates a Topological Transition that Drives Fiber Formation. Cell 111, 543– 551 (2002).

60. Shoji, M., Shibata, S., Sueyoshi, T., Naito, M. & Nakayama, K. Biogenesis of Type V pili. Microbiol. Immunol. 64, 643–656 (2020).

61. Böhning, J. et al. Molecular architecture of the TasA biofilm scaffold in Bacillus subtilis. bioRxiv 2022.03.14.484220 (2022) doi:10.1101/2022.03.14.484220.

62. Gambelli, L. et al. Architecture and modular assembly of Sulfolobus S-layers revealed by electron cryotomography. Proc. Natl. Acad. Sci. U. S. A. 116, 25278–25286 (2019).

63. Tsai, C.-L. et al. The structure of the periplasmic FlaG–FlaF complex and its essential role for archaellar swimming motility. Nat. Microbiol. 5, 216–225 (2020).

64. Tang, G. et al. EMAN2: An extensible image processing suite for electron microscopy. J. Struct. Biol. 157, 38–46 (2007).

65. Zivanov, J. et al. New tools for automated high-resolution cryo-EM structure determination in RELION-3. Elife 7, (2018).

66. Pettersen, E. F. et al. UCSF Chimera - A visualization system for exploratory research and analysis. J. Comput. Chem. 25, 1605–1612 (2004).

67. Sanchez-Garcia, R. et al. DeepEMhancer: a deep learning solution for cryo-EM volume post-processing. *Commun*. Biol. 4, (2021).

68. Goddard, T. D. et al. UCSF ChimeraX: Meeting modern challenges in visualization and analysis. Protein Sci. 27, 14–25 (2018).

69. Emsley, P. & Cowtan, K. Coot: Model-building tools for molecular graphics. Acta Crystallogr. Sect. D Biol. Crystallogr. 60, 2126–2132 (2004).

70. Altschul, S. F. et al. Gapped BLAST and PSI-BLAST: A new generation of protein database search programs. Nucleic Acids Res. 25, 3389–3402 (1997).

71. McGinnis, S. & Madden, T. L. BLAST: At the core of a powerful and diverse set of sequence analysis tools. Nucleic Acids Res. 32, (2004).

72. Cramer, P. AlphaFold2 and the future of structural biology. Nat. Struct. Mol. Biol. 28, 704–705 (2021).

73. Vagin, A. & Teplyakov, A. MOLREP: An Automated Program for Molecular Replacement. J. Appl. Crystallogr. 30, 1022–1025 (1997).

74. . The CCP4 suite: Programs for protein crystallography. Acta Crystallogr. Sect. D Biol. Crystallogr. 50, 760–763 (1994).

75. Lebedev, A. A. et al. JLigand: A graphical tool for the CCP4 template-restraint library. Acta Crystallogr. Sect. D Biol. Crystallogr. 68, 431–440 (2012).

76. Murshudov, G. N. et al. REFMAC5 for the refinement of macromolecular crystal structures. Acta Crystallogr. Sect. D Biol. Crystallogr. 67, 355–367 (2011).

77. Kanehisa, M., Furumichi, M., Tanabe, M., Sato, Y. & Morishima, K. KEGG: New perspectives on genomes, pathways, diseases and drugs. Nucleic Acids Res. 45, D353– D361 (2017).

78. Oberto, J. SyntTax: A web server linking synteny to prokaryotic taxonomy. BMC Bioinformatics 14, (2013).

79. Mirdita, M. et al. ColabFold - Making protein folding accessible to all. bioRxiv 2021.08.15.456425 (2022) doi:10.1101/2021.08.15.456425.

80. Krissinel, E. & Henrick, K. Secondary-structure matching (SSM), a new tool for fast protein structure alignment in three dimensions. Acta Crystallogr. Sect. D Biol. Crystallogr. 60, 2256–2268 (2004).

81. Sievers, F. et al. Fast, scalable generation of high-quality protein multiple sequence alignments using Clustal Omega. Mol. Syst. Biol. 7, (2011).

82. Afonine, P. V. et al. Towards automated crystallographic structure refinement with phenix.refine. Acta Crystallogr. Sect. D Biol. Crystallogr. 68, 352–367 (2012).

